# GH25 lysozyme mediates tripartite interkingdom interactions and microbial competition on the plant leaf surface

**DOI:** 10.1101/2025.04.04.647216

**Authors:** Zarah Sorger, Priyamedha Sengupta, Klara Beier-Heuchert, Jaqueline Bautor, Jane E. Parker, Eric Kemen, Gunther Doehlemann

**Author notes:** Contributed equally to the work (Zarah Sorger, Priyamedha Sengupta).

## Abstract

Microbial communities inhabiting plants have emerged as crucial factors in regulating plant health and defense against disease-causing pathogens. The basidiomycete yeast *Moesziomyces bullatus* ex. *Albugo* on *Arabidopsis* (*MbA)* releases Glycoside Hydrolase 25 (GH25) protein which regulates the leaf microbiome by antagonizing an oomycete *A. laibachii* biotrophic pathogen MbA. Application of both *MbA* and GH25 protein rescued fresh shoot weight of *A. thaliana* upon *A. laibachii* infection, showing its potential in plant protection. Tripartite interaction assays did not reveal antagonistic activity of GH25 towards other plant pathogenic oomycetes or fungi besides *A. laibachii*. We identified a core set of bacteria are closely associated with *A. laibachii* and established that GH25 inhibits members of this core group. Among *A. laibachii-*associated bacteria that were inhibited by GH25, *Curtobacterium sp*. could override the inhibition of *A. laibachii* by *MbA*. We describe a tripartite antagonistic interaction in which bacterium and oomycete protect each other from growth inhibition by *MbA. Curtobacterium sp*., in turn, exhibits specific inhibition of *A. laibachii*-associated bacteria that are not targeted by *MbA* but themselves antagonize *A. laibachii*. Our study reveals an inter-kingdom interaction network in which a GH25 lysozyme shapes the antagonistic relationship between yeast, a pathogenic oomycete and an oomycete-associated bacterium.

## Introduction

Pathogenic microorganisms such as bacteria, fungi, oomycetes and viruses cause various plant diseases. In particular, ooomycete and fungal pathogens infect various agronomically important crops; thereby, posing a threat to global food security. For example, the oomycete *Phytophthora infestans* caused the Irish famine (1845) through potato late blight disease, while other species of *Phytophthora*, namely *P. ramorum, P. capsici* and *P. sojae* are known to infect oak trees, solanaceous crops, and soybean, respectively (Kamoun et al., 2015). *Botrytis cinerea*, a fungal pathogen with a broad host range, can cause severe economic loses with the disease control being heavily dependent on fungicide application (De Angelis et al., 2022; Weiberg et al., 2013). In addition, climate change is predicted to exacerbate plant disease outbreaks through the emergence and evolution of pathogenic strains, potentially affecting both natural biodiversity and agricultural systems (Singh et al., 2023).

Microbial antagonism or biological control of pathogens by the action of beneficial microorganisms has emerged as a more environmentally sustainable approach towards plant protection. For example, biological control of *Phytophthora* has been reported by inhibitory activities of potato associated bacteria (De Vrieze et al., 2019), or by volatiles secreted from bacteria (Gfeller et al., 2022) and fungi (Oubaha et al., 2021). Therefore, a thorough knowledge on the mechanism of microbial antagonism is necessary to attain successful crop protection. While several modes of action for microbial antagonists have been described, such as triggering plant immune response, hyper-parasitism on pathogens or secretion of secondary metabolites (Köhl et al., 2019), competition by associated microbiota may be a crucial criterion for effective pathogen control and restoration of plant health (Cernava, 2024). Among such interactions, *Albugo laibachii*, the causal agent of white rust, has emerged as a microbial hub that promotes disease-associated microbiota in the Arabidopsis phyllosphere, in contrast to the health-associated communities found in uninfected plants (Mahmoudi et al., 2024).

Microbiota inhabiting plants secrete a range of glycoside hydrolases (GH) which aid nutrient acquisition and competition with other microbes, or can be virulence factors to colonize a host (Bradley et al., 2022). Antagonistic yeasts such as Trichoderma were reported to produce a variety of hydrolytic enzymes to antagonize fungal pathogens in crop plants (reviewed by Freimoser et al., 2019). For example, chitinase from *T. asperellum* PQ34 inhibited growth of fungal pathogens *Sclerotium rolfsii* and *Colletotrichum* (Bradley et al., 2022; Loc et al., 2020). Additionally, cellulose encoding genes from *T. harzianum* were found to elicit plant immunity against pathogen *F. graminearum* by upregulating the production of DIMBOA and other defense related genes in maize roots (Saravanakumar et al., 2018).

Glycoside hydrolases are one of the largest and most diverse groups among hydrolytic enzymes with 172 families and 18 different GH clans (Henrissat and Davies, 1997). In phytopathogenic fungi, GH families have been characterized mostly with respect to their suppression of plant immunity and pathogenic virulence. For example, secreted GHs from oomycete *P. sojae* and fungi *Verticillium dahliae, M. oryzae* and *F. oxysporum* can act as microbe-associated molecular patterns (MAMPs) (Gui et al., 2017; Ma et al., 2015; Zhang et al., 2021) or release damage associated molecular patterns (DAMPs) leading to the activation of pattern triggered immunity (PTI) (Bradley et al., 2022). In contrast, deletion of cell wall hydrolyzing enzymes encoded by GH families in *B. cinerea, M. oryzae* and *A. alternata* impairs fungal virulence (Brito et al., 2007; Ma et al., 2015; Yu et al., 2018; Zhang et al., 2021).

Glycoside hydrolases can also be involved in microbial antagonism of pathogens. For example, deletion of the GH18 chitinase gene in the fungus *Clonostachys rosea* reduced inhibitory activity against *B. cinerea* and *Rhizoctonia solani*, although biocontrol of *B. cinerea* was not compromised (Bradley et al., 2022; Tzelepis et al., 2015). More recently, Glycoside Hydrolase 30 (GH30) produced by the bacterium *Bacillus paralicheniformis* triggered plant defense responses via programmed cell death and restricted phytopathogens such as *Sclerotinia sclerotiorum, P. capisici* and tobacco mosaic virus (Yu et al., 2024). In addition, GH19 chitinase from *Chitinilyticum aquatile* CSC-1 was recombinantly expressed in *E. coli* and displayed inhibition of fungal growth in several *Fusarium* species, suggesting its role in biological control of phytopathogenic fungi (Yang et al., 2024).

In previous work (Eitzen et al., 2021) we described how the basidiomycete yeast, *Moesziomyces bullatus* ex. *Albugo* on *Arabidopsis* (*MbA)* regulates the *Arabidopsis thaliana* phyllosphere microbiota by inhibiting the white rust pathogen *Albugo laibachii* through a secreted Glycoside Hydrolase 25 (GH25) protein. The GH25 family of hydrolases, characterized by a DXE motif active site, cleaves the β-1,4-glycosidic linkage between N-acetylmuramic acid (NAM) and N-acetylglucosamine (NAG) in the bacterial peptidoglycan (Rau et al., 2001). However, the biological function of GH25 in fungi remains poorly understood.

In this study, we functionally investigated how GH25 antagonizes *A. laibachii*. We examined the effect of *MbA* and *MbA*_GH25 against other plant pathogens and found that GH25 acts specifically against *A. laibachii* through an indirect mechanism involving an associated bacterium of the genus *Curtobacterium*. We extended our analysis by functionally characterizing GH25 orthologs from the plant pathogenic fungi *Ustilago maydis* and *Rhizoctonia solani* in the antagonism towards *A. laibachii* and suggest a possible mechanism of evolutionary adaptation of GH25 specificity.

## Results

### *MbA* limits growth of *Albugo laibachii* through GH25 activity

In a previous study (Eitzen et al., 2021), we explored how a basidiomycete yeast, *Moesziomyces bullatus* ex *Albugo* on *Arabidopsis* (*MbA*) antagonizes the *oomycete Albugo laibachii* in the *A. thaliana* phyllosphere by secreting Glycoside Hydrolase 25. Recombinantly produced *MbA*_GH25 also reduced *A. laibachii* growth *in-planta*, which revealed the importance of GH25 in the antagonism of this oomycete pathogen.

In this study, we aimed to clarify how GH25 mediates the observed antagonism between *MbA* and *A. laibachii*. We first quantified *A. laibachii* infection symptoms in 3-week-old *A. thaliana* seedlings grown in the gnotobiotic plate system with and without application of *MbA* (**Fig. 1A**). Although *MbA* significantly reduced the white rust infection symptom at 14 dpi, for a more quantitative assessment, the relative fungal biomass *A. laibachii* at 10 dpi was analyzed using the oomycete internal transcribed spacer (ITS) 5.8s sequence normalized to the *A. thaliana* housekeeping gene ef1-α. We observed that *A. laibachii* biomass was also significantly reduced in the presence of *MbA* (**Fig. 1B**). Additionally, a protective effect of *MbA* on the plant was evaluated by measuring the fresh shoot weight of *A. thaliana* seedlings following different treatments (**Fig. 1C**). We observed that the reduced fresh shoot weight during oomycete infection could be reversed by pre-treating the plants with *MbA* or by adding recombinantly produced *MbA*_GH25 to the yeast deletion strain *MbA*Δgh25 (**Fig. 1C**).

**Figure 1:**
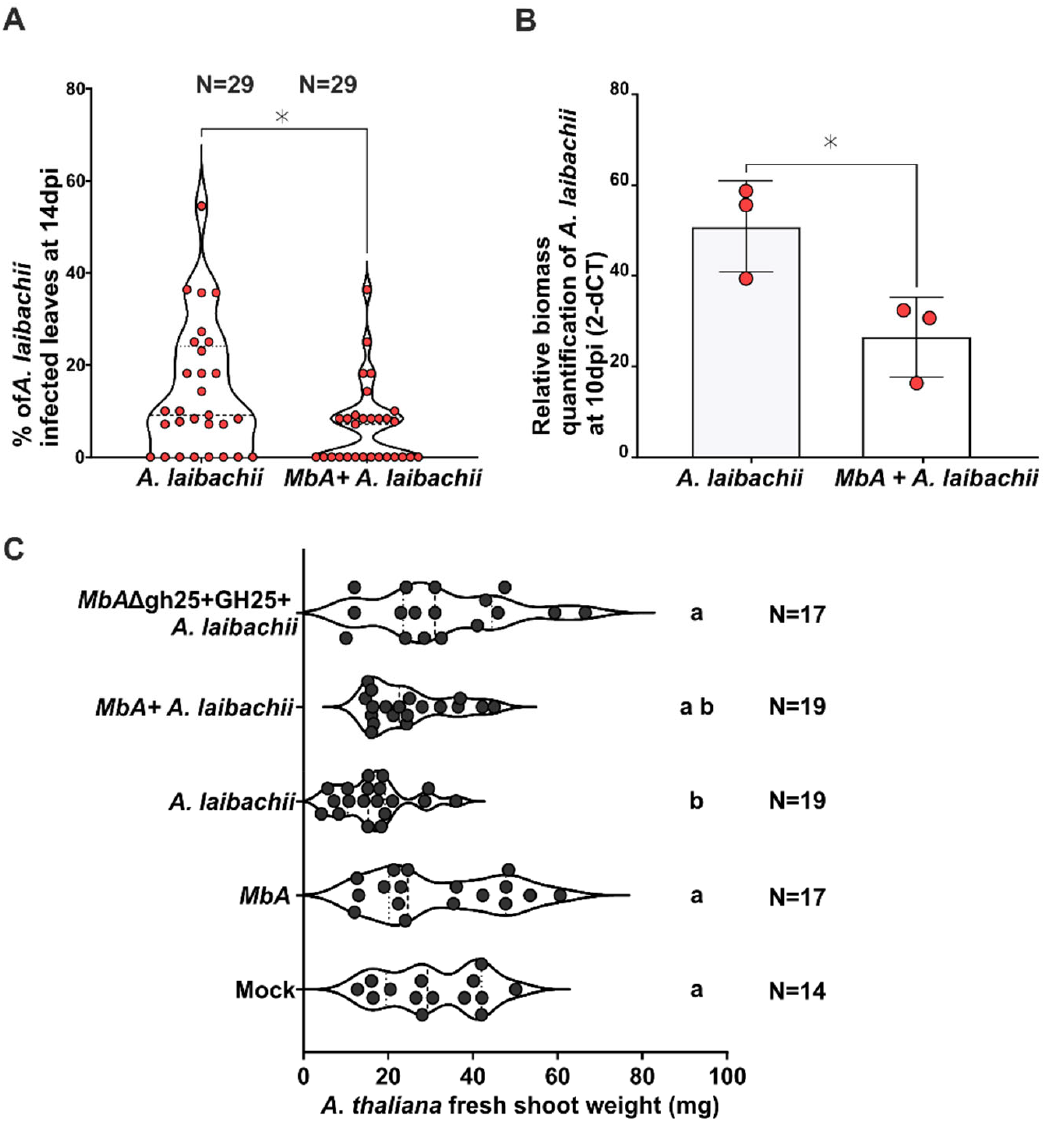
*MbA* restricts *A. laibachii* growth and recovers *A. thaliana* fresh shoot weight during infection. **A)** Addition of *MbA* reduces *A. laibachii* induced leaf disease symptoms of *A. thaliana* at 14 dpi. The number of infected plants (n) were scored across 3 biological replicates (unpaired t-test, p value =0.02). **B)** Relative quantification of *A. laibachii* biomass in response to *MbA* treatment by qPCR. The oomycete internal transcribed spacer (ITS) 5.8 s was normalized to *A. thaliana* EF1-α gene to quantify the amount of *A. laibachii* DNA in the samples at 10 dpi (unpaired t-test, p value =0.03). Error bars indicate SD. **C)** Fresh shoot weight measured at 11dpi showed *A. thaliana* seedlings to be reduced in growth upon infection with *A. laibachii* compared to mock treated seedlings. Addition of purified *MbA*_GH25 protein to the gh25 deletion mutant of *MbA* (*MbA*Δ*gh25*) led to a higher recovery of fresh shoot weight in *A. laibachii* infected seedlings. The number of seedlings (N) were measured across 3 biological replicates. One-way ANOVA and Tukey’ HSD (multiple comparisons of means; 95% family-wise confidence level) was performed to find significant difference between treatments. Representative image of *A. thaliana* seedlings is added next to each treatment.

Since *MbA*_GH25 plays a key role in oomycete inhibition, we analyzed the evolutionary conservation of GH25 in the fungal kingdom. Previously, we showed that the DXE active site motif is highly conserved in the GH25 amino acid sequence of different fungal groups such as Basidiomycetes, Ascomycetes and Chytrids (Eitzen et al., 2021). A newly constructed phylogenetic tree including various fungal species showed that the GH25 of Ustilaginales, including *MbA* and *U. maydis*, clustered together with a high degree of sequence similarity. However, evolutionarily more diverse ascomycete fungi, including several plant pathogens, also have significantly conserved GH25 orthologs (**Fig. 2A**). We recombinantly expressed orthologues from *MbA, U. maydis* and *Rhizoctonia solani* in FB1 strain of *U. maydis*. The recombinant strains were inoculated onto *A. thaliana* seedlings prior to *A. laibachii* infection and infection symptoms on leaves were scored at 11dpi. FB1 strains overexpressing active GH25s from *MbA* or *U. maydis* significantly inhibited *A. laibachii* infection *in-planta*, as opposed to GH25 from *R. solani* (**Fig. 2B**). Therefore, inhibitory activities of GH25 from *MbA* and *U. maydis* are functionally conserved compared to GH25 from the more distantly related fungal species *R. solani*.

**Figure 2:**
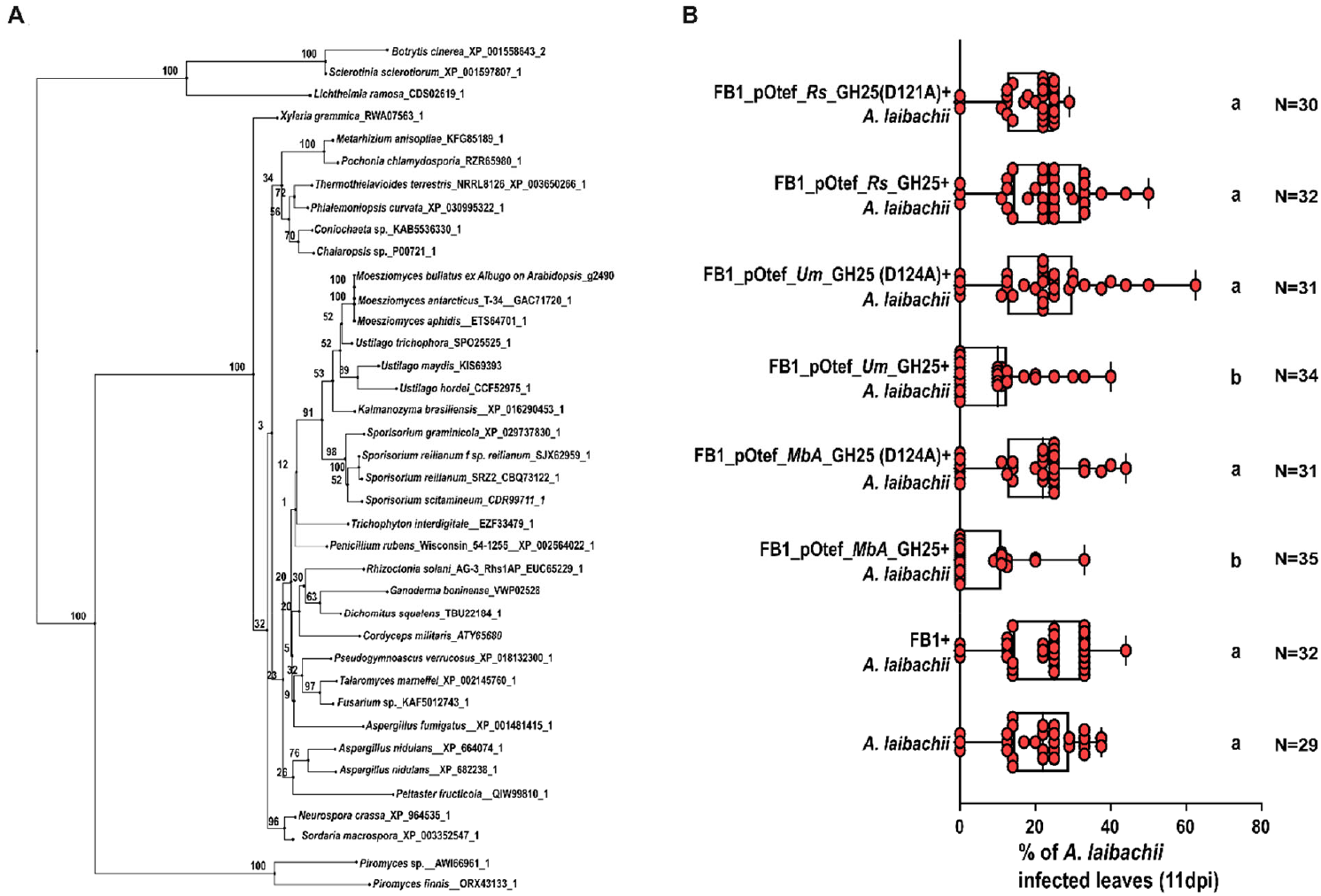
Overexpression of GH25 orthologues from *MbA* and *U. maydis* restricts *A. laibachii* growth compared to GH25 from distantly related *R. solani*: **A)** Phylogenetic tree of GH25 orthologues. Protein sequences were aligned using MAFFT version 7 multiple alignment program for amino acid of nucleotide sequences with default parameters. Phylogenetic tree construction was based on the alignment using Phylio.io. **B)** Infection assay of *A. laibachii* on *A. thaliana* with pre-treatment of *U. maydis* wild type strain FB1 and recombinant FB1 strains overexpressing GH25 orthologues from *MbA, U. maydis* (*Um*) and *R. solani* (*Rs*) (FB1_pOtef_\**MbA/Um/Rs*\*_GH25) in active or mutant version (D124A). % of *A. laibachii* infected leaves were scored at 11 dpi across 3 biological replicates in non-sterile condition. One-way ANOVA and Tukey’s multiple comparisons test (alpha 0.05) was performed to find significant difference between treatments. N=number of seedlings analyzed.

### No evidence of plant immunity activation or direct inhibition of oomycete and fungal plant pathogens by *MbA* GH25

To elucidate the mechanism of the observed Mba antagonism towards *A. laibachii*, we explored whether *MbA*_GH25 impacts plant immunity. To this end, 2.5-week-old *A. thaliana* seedlings were treated with purified recombinant *MbA*_GH25, *MbA*_GH25 (D124A) or heat-inactivated (Hi) *MbA*_GH25 proteins as a negative control. The broadly recognized bacterial PAMP flagellin 22 (flg22), was used as a defense-inducing positive control in the elicitor assay. Thirty minutes after treatments, seedlings were harvested to check for activation of several defense marker genes. Flg22 treatment induced the expression of defense genes *WRKY53, WRKY33, WRKY30*, and *FRK1* compared to other treatments (**Fig. S1**). No difference in defense gene expression levels was detected between active and inactive (mutated and heat-inactivated) versions of the *MbA*_GH25 protein (**Fig. S1**). Therefore, we concluded that the enzymatic activity of *MbA*_GH25 likely does not induce plant defense in *A. thaliana*.

Since the *MbA*_GH25 mediated antagonism of *A. laibachii* does not depend on an induction of the plant PTI system, we tested whether *MbA*_GH25 directly targets the oomycete. The cell walls of oomycetes are mainly composed of β-1,3, and β-1,6 glucans (Aronson et al., 1967), with varying levels of N-acetyl Glucosamine (NAG)-the building blocks of bacterial peptidoglycan (Mélida et al., 2013). Therefore, to determine whether *MbA_*GH25 directly targets *A. laibachii* cell wall we used commercially available Laminarin (seaweed polysaccharide with ß-1,3 linked glucan bonds and ß-1,6 linked side chains) as a substrate to monitor for release of oligosaccharides upon addition of purified *MbA*_GH25 protein via Thin Layer Chromatography. This approach showed no activity of GH25 on Laminarin (**Fig. S2A**). Additionally, we isolated *A. laibachii* cell walls from harvested zoospores (Mélida et al., 2013), and looked for activation of *MbA* gh25 gene expression. However, no significant induction of the *gh25* expression was detected in growing *MbA* cultures (**Fig. S2B**).

To determine whether the antagonistic activity of *MbA*/GH25 is specific to *A. laibachii*, we tested for possible interactions with the oomycete pathogen *Hyaloperenospora arabidopsidis* (*Hpa;* biotrophic pathogen, adapted to *A. thaliana*) and *Phytophtora infestans* (hemibiotrophic pathogen of solanaceous plants, not adapted to *A. thaliana*) as well as the fungal pathogen *Botrytis cinerea* (nectrotrophic pathogen, generalist). To test for *Mba*/GH25 effects on *Hpa* infection, 2.5-week-old *A. thaliana* seedlings (Col-0 and Col-0 *eds1-12* (Ordon et al., 2017)) were pre-treated with a growing culture (OD_600nm_=0.8-1) of *MbA* and *MbA*Δgh25, followed by spray inoculation of *Hpa* (15*10^4^ spores/ml) 2 days later. Purified *MbA* GH25 or *MbA* GH25(D124A) was mixed directly with *Hpa* spores (6µM conc.) and sprayed on seedlings. No significant differences in *Hpa* sporulation (release of *Hpa* spores per gram of *A. thaliana* at 5 dpi) were observed among the different treatments in both *A. thaliana* wild-type Col-0 and the *Hpa*-hypersusceptible mutant *eds1-12* (**Fig. 3A**). *Hpa* sporulation was significantly higher in the eds1-12 mutant line compared to Col-0, indicating that *Hpa* infection conditions on *A. thaliana* were suitable. (**Fig. 3A**). Furthermore, relative gh25 expression levels were quantified in *MbA* upon interaction with *Hpa* on *A. thaliana* and no significant change in gh25 expression was observed (**Fig. S3A**). Thus, the presence of the basidiomycete yeast or its active and inactive hydrolase did not impact leaf infection *Hpa*.

**Figure 3:**
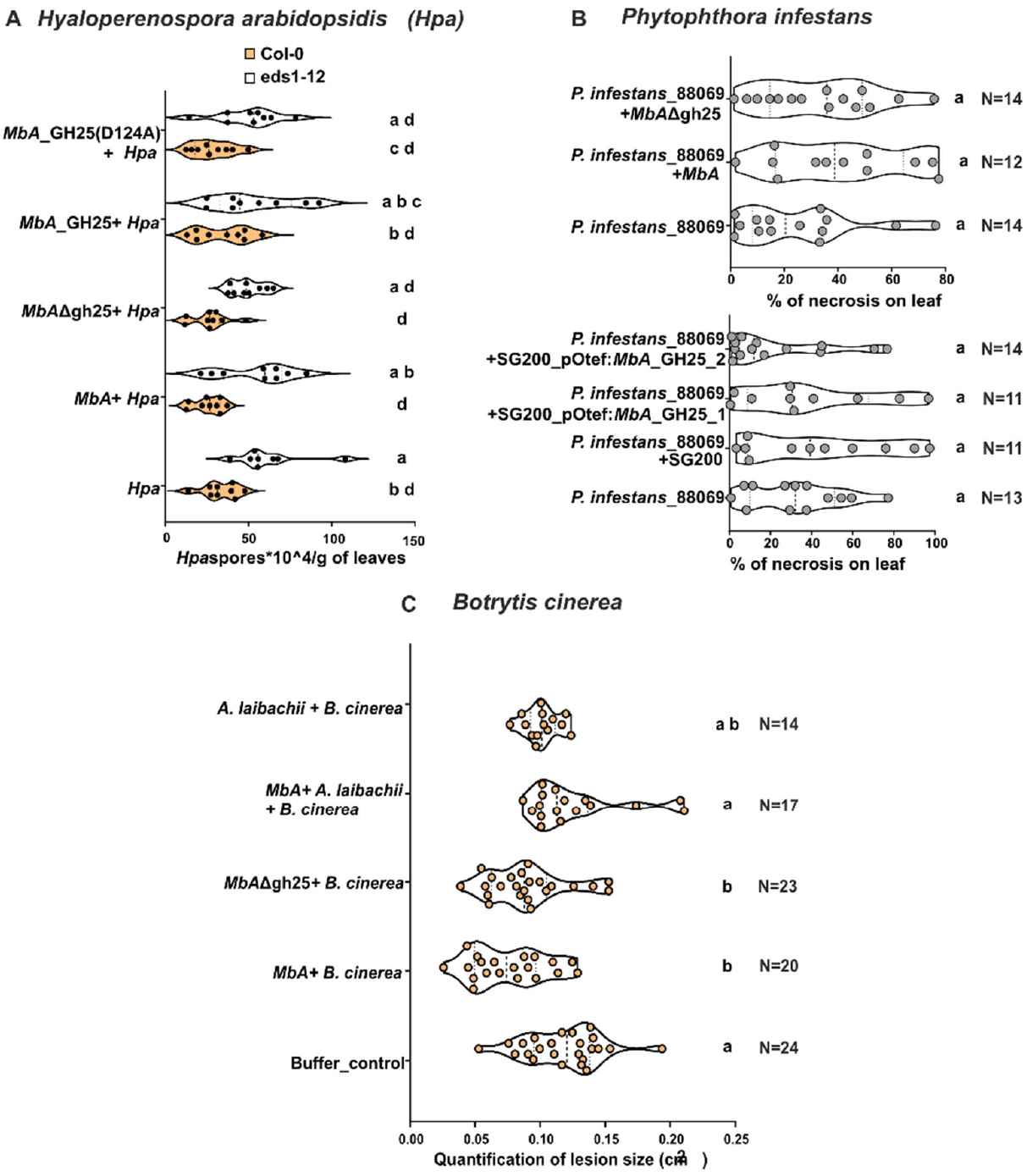
GH25-mediated inhibition of *A. laibachii* is not conserved against other pathogenic oomycetes and fungi. **A**) *MbA* and purified *MbA*_GH25 do not affect *Hpa* infection in *A. thaliana* Col-0 and eds1-12 mutant (Col-0 background). Experiments were conducted in 3 biological replicates (consisting of 3 technical replicates). Quantification of *Hpa* spores *10^4^ / g of leaves was performed at 5 dpi. **B)** Interaction of *P. infestans* with *MbA* strains and *U. maydis* strains: Quantification of necrotic area on leaf surface was performed by ImageJ and percentage of necrosis calculated in 3 biological replicates (N=number of leaves analyzed). **C)** *In planta* droplet infection assay was performed by dropping 5 µl of spore suspension in detached A. thaliana leaves, which were pretreated with mock (water), *MbA, MbA*)GH25, *MbA* + *A. laibachii* or *A. laibachii*. Lesion size quantification was performed using ImageJ across 3 biological replicates (N= number of lesions analyzed). One-way ANOVA and Tukey’s multiple comparisons test (alpha 0.05) was performed to find significant difference between treatments.

*In-planta* interaction assays of *P. infestans* with *MbA* were performed using detached leaves of 5-week-old *N. benthamiana*. We observed that necrotic lesions caused by *P. infestans* were not reduced upon its co-inoculation in the presence of *MbA* and *MbA*Δgh25 (**Fig. 3B**). In addition, the *MbA* gh25 gene was recombinantly expressed in the smut pathogen *U. maydis* (SG200_pOtef:*MbA*GH25) and tested for interaction with *P. infestans* in the detached leaf assay. However, co-inoculation of the recombinant strain SG200_pOtef:*MbA*GH25 did not limit the development of necrotic lesions (**Fig. 3B**).

Confrontation assays in axenic culture between *MbA* strains and *P. infestans* were carried out on Rye-Sucrose Agar (RSA) plates. However, although no zone of inhibition was observed between the two interacting microorganisms, *P. infestans* was restricted from growing over the area already colonized by yeast strains (**Fig. S3B**). To investigate the inhibitory effect of *MbA* on a necrotrophic fungus, *Botrytis cinerea* was tested by using both *in-vitro* and *in-planta* assays. During the axenic confrontation assay, no inhibition of *B. cinerea* could be observed upon confrontation with *MbA, MbA*Δgh25 and purified *MbA*_GH25 protein (**Fig. S3C**). However, the lesion size of *B. cinerea* was significantly reduced after pretreatment of *A. thaliana* with both *MbA* and *MbA*Δgh25, indicating an inhibition of *B. cinerea* by *MbA* that is independent of GH25 activity (**Fig. 3C**). Thus, the observed antagonistic activity of *MbA* via GH25 appears to specifically act against *A. laibachii*.

In summary, we found no evidence for a GH25 stimulating activity on plant immunity or a GH25 direct antimicrobial activity against any of the tested eukaroytic plant pathogens. Therefore, we hypothesized that GH25-mediated inhibition of *A. laibachii* is mediated by an indirect interaction involving additional microbes.

### Bacteria associated with *Albugo laibachii* are targeted by GH25

Recently, Goossens et al. (2023) investigated the presence of several bacterial isolates closely associated with *H. arabidopsidis* on *A. thaliana*. The bacterial population thrived during *Hpa* infection and was actively recruited by the plant to limit disease progression (Goossens et al., 2023). Here, we tested whether *A. laibachii* could be associated with a bacterial community. In previous gnotobiotic *A. laibachii* infection assays, oomycete zoospores were treated with an antibiotic cocktail (streptomycin, kanamycin, rifampicin, and geneticin) prior to inoculation of *A. thaliana* seedlings. To determine whether any bacterial isolates remained after the antibiotic treatment, we harvested zoospores of *A. labachii* NC14 strain and plated them on King’s B medium after treatment with the antibiotic cocktail. This treatment resulted in the isolation of a set of *A. laibachii-*associated bacteria, comprising mainly *Pseudomonas* sp., *Microbacterium* sp. and *Curtobacterium* sp. (**Table S2**).

We then tested the individual *A. laibachii* associated bacterial strains in one-to-one interactions with *MbA*, purified *MbA*_GH25 protein or a recombinant *U. maydis* strain constitutively expressing *MbA*_GH25 (**Fig. 4, S4**). *MbA* inhibited two of the *Pseudomonas* strains, while the recombinant *MbA*_GH25 strain inhibited two of the *Curtobacterium* strains, although halo formation was more pronounced in the case of *MbA* (**Fig. S4**). Purified *MbA*_GH25 inhibited both *Stapylococcus* sp. and *Curtobacterium* isolates. In contrast, the catalytically inactive *MbA*_GH25 (D124A) protein did not cause any inhibition, indicating that the antibacterial activity is dependent on GH25 enzymatic activity (**Fig. 4A-D**). Thus, *A. laibachii* is associated with a community of bacteria of which *Stapylococcus* sp. and *Curtobacterium* sp. are inhibited by *MbA* / GH25 activity. Next, we tested whether these microbial interactions are linked with the antagonistic interplay of *MbA* and *A. laibachii* on the leaf surface.

**Figure 4:**
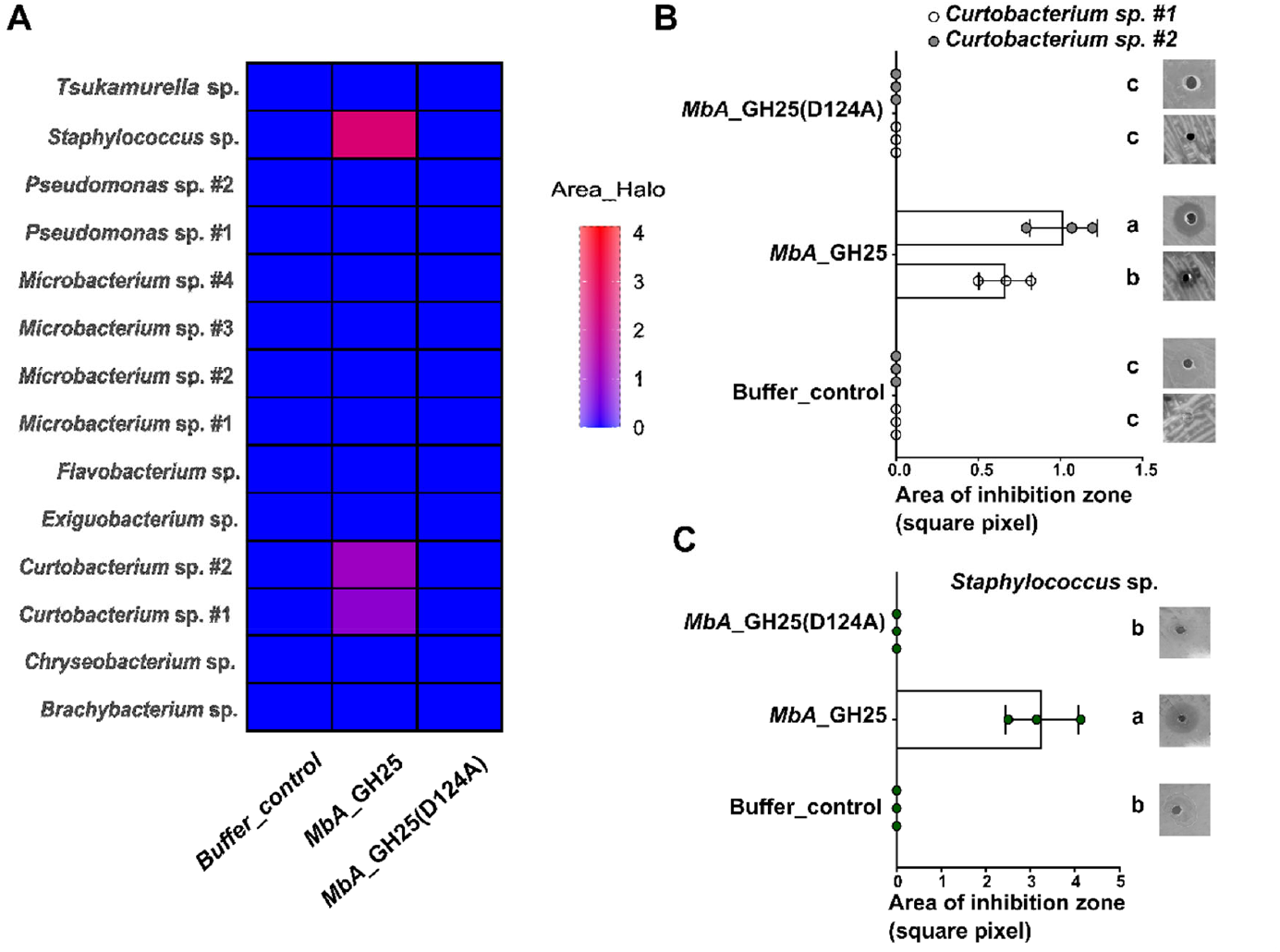
*Albugo laibachii* associated bacteria are targeted by *MbA*_GH25. **A)** Heatmap showing overview of the confrontation assay between *A. laibachii* associated bacteria and purified recombinant *MbA*_GH25/*MbA*_GH25(D124A) protein across 3 independent replicates. *Curtobacterium* sp. (#1 and #2) and *Staphylococcus* sp. were inhibited by *MbA*_GH25 active protein. **B)** Zone of inhibition of both isolates of *Curtobacterium* sp. and *Staphylococcus* sp. by purified *MbA*_GH25 was analyzed compared to buffer control and mutated *MbA*_GH25 (D124A) across 3 independent replicates. Representative image of bacterial inhibition added next to respective treatments as indicated by the presence or absence of halo formation surrounding the agar well, 3 days after application of recombinant protein (0.8ug/ul). Error bars show SD. One-way ANOVA and Tukey’s multiple comparisons test (alpha 0.05) was performed to find significant difference between treatments.

### An antagonistic interplay between bacteria, an oomycete and a fungal yeast

To elucidate an eventual biological role of the observed microbial interplay, the tripartite interactions of *MbA, A. laibachii* and either *Curtobacterium sp*. or *Staphylococcus sp*. on *Arabidopsis* leaves were investigated. While *Pseudomonas sp*. and *Staphylococcus sp*. had no, or marginal negative impact on *Albugo laibachii*, oomycete infection was strongly influenced by presence of *Curtobacterium sp*. (**Fig. 5A-C**). *Staphylococcus* sp. inhibited *A. laibachii* infection both in the presence and absence of *MbA*, similar to *Curtobacterium sp. #1* (**Fig. 5C**). In contrast, *Pseudomonas* sp. had no inhibitory effect on *A. laibachii* infection and partially suppressed the inhibition of *A. laibachii* by *MbA* (**Fig. 5C**), suggesting that *Pseudomonas* sp. positively affects *A. laibachii* colonization. Interestingly, the two strains of *Curtobacterium* sp. showed contrasting behavior in the tripartite interaction: *Curtobacterium sp. #1* reduced *A. laibachii* infection similarly to *MbA* (**Fig. 5A**). In contrast, *Curtobacterium sp. #2* did not significantly affect *A. laibachii* infection. Moreover, co-inoculation almost completely restored *A. laibachii* infection in presence of *MbA;* i.e. *Curtobacterium sp. #2* protected *A. laibachii* from antagonism by *MbA* (**Fig. 5B**).

**Figure 5:**
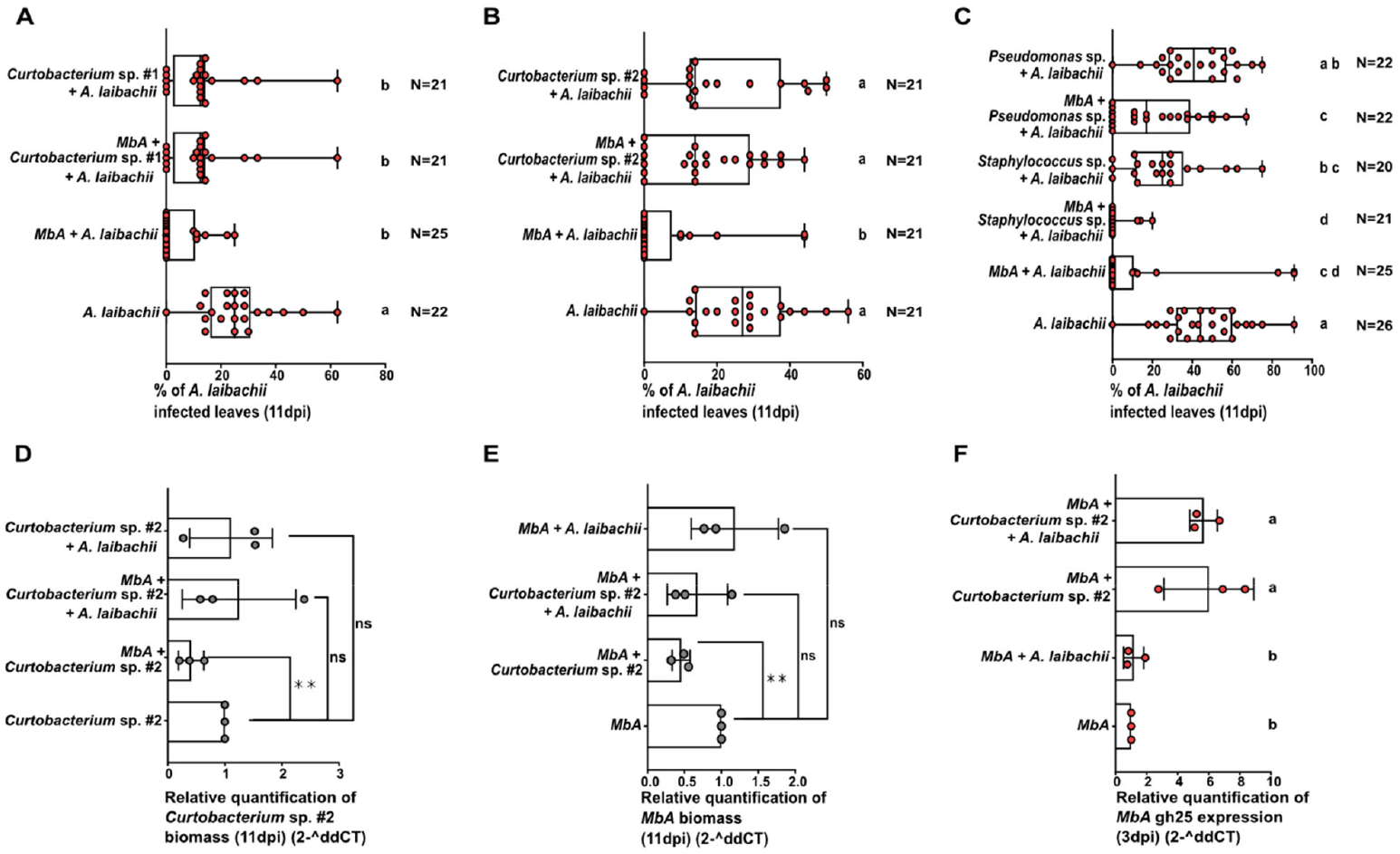
*Curtobacterium* sp. #2 suppresses *MbA* inhibition of *A. laibachii*. **A-C)** Infection assay of *A. laibachii* on *A. thaliana* with pre-treatment of *MbA* and different bacteria; **A)** *Curtobacterium* sp. #1, **B)** *Curtobacterium* sp. #2, **C)** *Staphylococcus* sp. and *Pseudomonas* sp. Pretreatment was performed 2 days prior to *A. laibachii* infection. Plants were scored at 11 dpi. One-way ANOVA and Tukey’s multiple comparisons test (alpha 0.05) was performed to find significant difference between treatments. **D)** Relative quantification of *Curtobacterium* sp. #2 biomass in response to *A. laibachii* and *MbA*. 23S rRNA specific region in *Curtobacterium* sp. was normalized to *A. thaliana* EF1-α to quantify the amount of *Curtobacterium* DNA in the samples at 11dpi. **E)** Relative quantification of *MbA* biomass in response to *A. laibachii* and *Curtobacterium* sp. GH25 (g2490) as a single-copy gene was normalized to *A. thaliana* EF1-α to quantify the amount of *MbA* DNA in the samples at 11dpi. Unpaired t-test carried out to find significant difference between treatments. Both *Curtobacterium* sp. #2 and *MbA* biomass was significantly reduced upon interaction with one another. **F)** Relative quantification of *MbA* gh25 expression in response to *A. laibachii* and *Curtobacterium* sp. #2, normalized to *MbA* housekeeping gene ppi at 3dpi. One-way ANOVA and Tukey’s multiple comparisons test (alpha 0.05) was performed to find significant difference between treatments.

Next, we performed a qPCR approach to quantify the growth of the interacting microbes on the plant surface. This revealed that *MbA* biomass was reduced in the presence of *Curtobacterium sp*.*2* (**Fig. 5E**). Also growth of *Curtobacterium sp. #2* was also significantly reduced in presence of *MbA* (**Fig. 5F**), reflecting a bidirectional antagonism between *Curtobacterium sp. #2* and *MbA*. Notably, the *MbA*-mediated inhibition of *Curtobacterium sp. #2* was rescued by the presence of *A. laibachii*, suggesting a protective effect of *A. laibachii* on *Curtobacterium sp. #2*. An interesting observation was made regarding the transcriptional induction of *MbA_gh25*. In our previous study (Eitzen et al., 2021), we had initially identified *gh25* as a candidate gene based on its transcriptional activation on the plant surface specifically in the presence of *A. laibachii*. Here we find that *gh25* expression in *MbA* was significantly upregulated in the presence of both *A. laibachii* and *Curtobacterium* sp. *#*2. However, also confrontation of *MbA* with *Curtobacterium* sp. *#*2 alone resulted in transcriptional activation of *gh25*. In contrast, antibiotic-treated *A. laibachii* alone did not induce *gh25* expression in *MbA* (**Fig. 5D**). We concluded from the above assays that it is not *A. laibachii* but its associated *Curtobacterium* sp. *#*2, which directly antagonizes *MbA*, that is the trigger of *gh25* expression.

Next we investigated whether the observed strain-specific inhibitory activity of GH25 was restricted to *MbA*_GH25 or might be a conserved property of GH25 proteins across clades. To test this, we performed confrontation assays with the *FB1* strains overexpressing GH25 orthologs derived from *MbA, U. maydis* or *R. solani* (see **Fig. 2A**) and *Curtobacterium sp. #2*. Consistent with GH25 antagonism of *A. laibachii* infection, GH25 from *MbA* and *U*.*maydis* inhibited *Curtobacterium sp. #2* (**Fig. S5A**). In contrast, GH25 from *R. solani* had no effect on *Curtobacterium sp. #2*, whereas a bacterial control strain (*Priestia* sp.) was inhibited by enzymatically active versions of all three GH25 orthologs (**Fig. S5B**). This result suggests an ortholog-dependent target specificity of GH25, which may reflect a host-specific adaptation. Finally, we tested whether the protective effect of *Curtobacterium* sp. #2 on *A. laibachii* can be attributed to its modulation of the leaf microbiome. We examined potential cross-inhibition within the microbial community by spreading one bacterium as a lawn and applying treatment bacteria to observe any resulting halo formation (**Fig. S6A**). This showed that *Curtobacterium* sp. #1 and #2, as well as *Pseudomonas* sp. #1 and #2, inhibited most of the associated bacteria that were not targeted by GH25 (**Fig. 6A**). Furthermore, when these bacteria were used as a lawn, the inhibited bacteria were unable to grow, suggesting strong inhibitory potential (**Fig. S6B**). Based on these findings, we hypothesized that *Curtobacterium* sp. and *Pseudomonas* sp. might promote *A. laibachii* growth through their inhibitory effects on other microbes. To test this, we conducted an infection assay using *Microbacterium* sp. #2 and *Chryseobacterium* sp., two bacteria that not inhibited by GH25 but are targeted by both *Pseudomonas* sp. and *Curtobacterium* sp. We found that both *Microbacterium* sp. #2 and *Chryseobacterium* sp. displayed an inhibitory effect on *A. laibachii* and were unable to alleviate inhibition of *A. laibachii* by MbA (**Fig. 6B**). In addition, they rescued the growth inhibition of *A. thaliana* caused by *A. laibachii* (**Fig. S6C**). Taken together, our findings reveal a tripartite, inter-kingdom interaction between bacteria, oomycete and yeast, in which a GH25 lysozyme shapes a highly specific antagonism between *MbA, A. laibachii* and *Curtobacterium sp*.. We propose that this interplay between organisms on the leaf enables a balanced competition and maintenance of microbial niches in the plant phyllosphere (**Fig. 7**).

**Figure 6:**
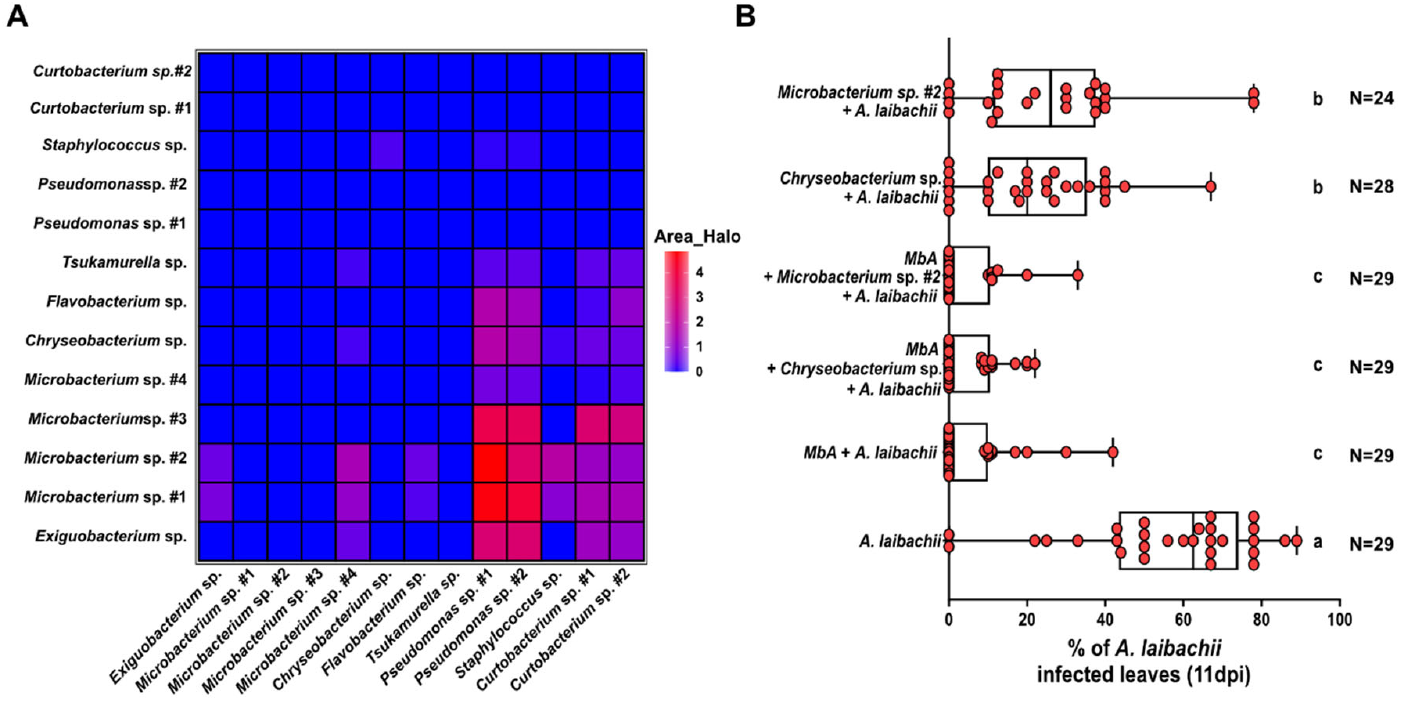
Cross-inhibition events within the associated bacterial community shape infection outcome. **A)** Inhibition assays between bacteria were conducted on square PD plates. 250 µl lawn bacterium was plated out and 5 µl competitor/treatment bacterium spotted on top. After 3-4 days halo area was measured and analyzed using ImageJ. 3-4 biological were performed for each bacterial pair tested. Mean of replicates is used for plotting. **B)** Infection assay of *A. laibachii* on *A. thaliana* with pre-treatment of MbA and *Microbacterium* sp._#2 and *Chryseobacterium* sp. Pretreatment was performed 2 days prior to *A. laibachii* infection. Plants were scored at 11 dpi. One-way ANOVA and Tukey’s multiple comparisons test (alpha 0.05) was performed to find significant difference between treatments.

**Figure 7:**
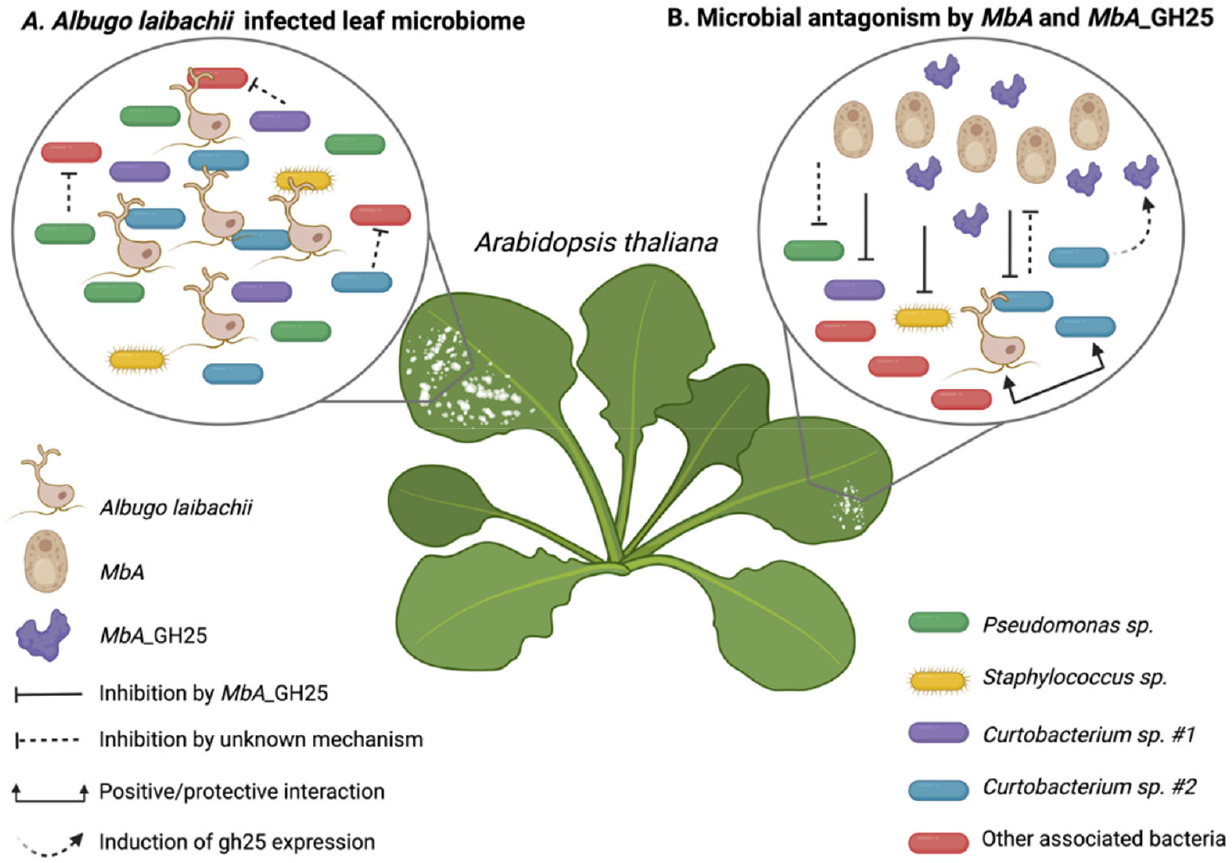
Model of the inter kingdom interactions shaped by *MbA*GH25. We depict how white rust infection in *A. thaliana*, caused by oomycete *A. laibachii* is inhibited by basidiomycete yeast *MbA* through antagonism of oomycete associated bacteria. **A)** *Albugo laibachii* is closely associated with a bacterial consortium in the phyllosphere of *A. thaliana*. Particularly, *Curtobacterium* sp. and *Staphylococcus* sp. are positively correlated with the host infection caused by *A. laibachii*. **B)** Basidiomycete yeast *MbA* produces Glycoside Hydrolase 25 protein to target *Curtobacterium* sp. #2, which leaves *A. laibachii* unprotected from competition with *MbA*, leading to a reduction of white rust infection in *A. thaliana*.

## Discussion

We previously described antagonism by the basidiomycete yeast *MbA* of an oomycete plant pathogen *A. laibachii* through an MbA-secreted GH25 lysozyme (Eitzen et al., 2021). Here, we demonstrate that the observed antagonism between *MbA* and *A. laibachii* is mediated through an *A. laibachii* associated *Curtobacterium* sp. in a strain-specific manner.

We tested two different modes of antagonism, including a direct action of the GH25 on the oomycete cell wall, as well as an indirect inhibition of infection via the activation of plant immune responses. Simultaneously, we found that *MbA* and Glycoside Hydrolase 25 were unable to antagonize oomycete pathogens *Hpa* and *P. infestans*. Although both *A. laibachii* and *Hpa* are foliar pathogens in natural *A. thaliana* populations, *Hpa* is less abundant and more affected by hormonal alterations in the host (Ruhe et al., 2016). For instance, while *A. laibachii* growth is unperturbed in ABA and SA mutant lines of *A. thaliana, Hpa* growth is restricted on JA accumulating ABA mutant line and increased on sid2-2 mutant, defective in Isochorismate Synthase 1 (precursor in SA signaling) (Ruhe et al., 2016). Nevertheless, *A. candida* is known to suppress broad spectrum host innate immunity and increase susceptibility of the plant to downey mildew infection (Cooper et al., 2008; Prince et al., 2017). Several reports have shown *Hpa* disease symptoms to be controlled by the presence of bacteria. Berendsen et al. (2018) reported three bacterial taxa (*Stenotrophomonas* sp., *Xanthomonas* sp., and *Microbacterium* sp. to become enriched in the *A. thaliana* rhizosphere upon infection with *Hpa*. More interestingly, in a study by Almario et al. (2022), although *Hpa* comprised the core taxa, the sampled leaves were asymptomatic due to the presence of plant beneficial bacterium such as *Sphingomonas* and *Variovorax*. Whereas, for *P. infestans*, a fungal endophyte, *Monosporacus* sp. inhibited *Phytophthora* in culture conditions and not *in planta* (de Vries et al., 2018). Since both *Hpa* and *A. laibachii* are obligate biotrophic pathogens, their survival depends on the living host and therefore maintaining a stable equilibrium in the host-microbial community is crucial (Agler et al., 2016; Ruhe et al., 2016).

Exploring microbiota associated with pathogens can offer insights into disease emergence and proliferation (Kong and Hong, 2016). For example, *A. laibachii* infections help *P. infestans* to colonize and sporulate on *Arabidopsis* (Belhaj et al., 2017), which could result from *Albugo* infections in Brassicaceae promoting a host jump by certain pathogens (Thines, 2014). *Albugo candida* has been shown to directly influence the plant microbiome by releasing proteins and peptides with antimicrobial activity into the apoplast (Gómez-Pérez et al., 2023). Moreover, microbiome analysis during the interaction between *A. thaliana* and *A. laibachii* revealed the formation of a disease associated microbial community (DCom) in infected plants, distinct from the health-associated microbial community (HCom) found in uninfected controls (Mahmoudi et al., 2024). Furthermore, co-inoculation with HCom members more effectively suppressed *A. laibachii* infection than co-inoculation with DCom-associated microbes (Mahmoudi et al., 2024).

Therefore, we were intrigued to explore how associated bacteria of *A. laibachii* might be a key component in the *MbA*_GH25 mediated inhibition of *A. laibachii*. To this end, we isolated 14 bacterial members from *A. laibachii* Nc14 spores and tested for interaction with *MbA* and *MbA*_GH25. During the confrontation assays, we found the *Pseudomonas* sp. strains to be inhibited by *MbA*, and not *MbA*_GH25, which indicates that the antimicrobial activity against *Pseudomonas* is independent of GH25 secretion. The *MbA* induced bacterial inhibition could be potentially caused by bio-surfactants or secondary metabolites as described previously for related yeasts (Morita et al., 2007). Interestingly, both *Curtobacterium sp*. isolates were inhibited by GH25 protein, as well as by recombinant *U. maydis* strains overexpressing GH25. However, only *Curtobacterium sp. #2* rescued the inhibition of *A. laibachii* by *MbA*, reflecting a high level of specificity. Moreover, both *Curtobacterium* sp. *#2* and *MbA* significantly antagonize each other *in-planta*, indicating a bi-directional negative interaction. Finally, *gh25* gene expression in *MbA* was upregulated in presence of *Curtobacterium sp. #2* rather than *A. laibachii*. Furthermore, we found *Curtobacterium sp*. and *Pseudomonas sp*. to be key microbes in shaping the microbiome associated with *A. laibachii*, by inhibiting most isolated associated bacteria that are not targeted by GH25. Taken together, our observations reveal a tripartite antagonism between *MbA, Curtobacterium* sp. *#2* and *A. laibachii*, which results in reshaping of the microbial community associated to *A. laibachii*. Together with results that demonstrate a preference of *A. labachii* towards DCom microbes (Mahmoudi et al., 2024) and the general role of *A. laibachii* as a hub microbe in shaping the microbial community (Agler et al., 2016) one could hypothezised that the association with *Curtobacterium sp. #2* aids *A. laibachii* in maintaining the stability of its niche. Unlike *Hpa*, which is associated with bacteria inhibiting the pathogens infection (Goossens et al., 2023), *A. laibachii* thus might have developed mechanisms to maintain a microbiome including protective microbes.

Simultaneously, we tested GH25 orthologues from *U. maydis* and *R. solani* in the antagonism towards A. *laibachii*. We found that GH25 from closely related basidiomycetes *MbA* and *U. maydis* were able to inhibit both *A. laibachii in-planta* and *Curtobacterium* sp. #2 during *in vitro* confrontation assays, as opposed to that from phylogenetically more distant plant pathogen *R. solani*. Therefore, these results indicate a host-specific adaptation of GH25 orthologues across the fungal kingdom and underline the importance of *Curtobacterium* sp. #2 inhibition for successful *A. laibachii* inhibition in-planta.

It has been shown that filamentous plant pathogens secrete effectors to modulate the plant microbiota (Flores-Nunez and Stukenbrock, 2024; Snelders et al., 2022, 2018). For example, the secretion of phytotoxic ribonucleases by fungal pathogens *Zymoseptoria tritici* (Kettles et al., 2018) and *U. maydis* (Ökmen et al., 2023)can modify the leaf microbiome of their respective hosts. In case of GH25 specificity, one could hypothesize that adaptation to competing bacteria in the leaf phyllosphere might contribute to niche adaptation in pathogenic fungi, but also increases fitness of beneficial fungi such as *MbA* to establish itself in an epiphytic lifestyle in association with highly competitive *A. laibachii*.

In conclusion, competition with associated microbes of plant pathogens can be a key component in effective disease control and plant health restoration (Cernava, 2024). Our study reveals a tripartite interkingdom interaction between the yeast *MbA*, the oomycete *A. laibachii*, and the bacterium *Curtobacterium* sp. *#2* on the plant surface. We show that *MbA* and *Curtobacterium* sp. *#2* exhibit mutual antagonism, with *A. laibachii* providing a protective effect on *Curtobacterium* sp. *#2* and vice versa. Our data further suggests a host-specific adaptation of GH25 orthologues in microbial competition and direct targeting of specific bacteria leading to a reshaping of the microbial community structure. Here, we propose a model depicting the cross-kingdom interaction between bacteria, oomycete and yeast orchestrated by GH25 (**Fig. 7**).

## Materials and methods

### Growth conditions for microbial strains

*MbA* and *U. maydis* strains were grown in liquid YEPS light medium at 22°C and 28°C respectively in a rotary shaker (200 rpm) and maintained on PD agar plates. *Albugo laibachii* associated bacterial strains (**Table S2**) were grown in King’s B liquid media at 22°C overnight in a rotary shaker (200 rpm). *Pichia pastoris* KM71H-OCH was used for recombinant protein expression as described in Eitzen et al. (2021).

### Plant infection assays

*Albugo laibachii* infections in *Arabidopsis thaliana* were performed as described in (Eitzen et al., 2021). *A. laibachii* infections of the orthologue experiment were performed in non-sterile conditions. Fresh shoot weight was taken at 11 dpi and roots were removed for the measurements. For *Hyaloperenospora arabidopsidis* (*Hpa*) infection assays, *Hpa* isolate Noco2 (van der Biezen et al., 2002)was sprayed onto 3-week-old *Arabidopsis thaliana* Col-0 and Col *eds1-12* mutant (Ordon et al., 2017) seedlings, two days after spraying with *MbA* strains and placed in a controlled environment under 10-h light/14-h dark regime at 22°C and 60% relative humidity. At 5dpi, plant fresh weight was determined and seedlings were resuspended in 5ml of ddH_2_O to release the conidiospores of *Hpa*. Conidiospores were counted under the light microscope using a Neubauer Chamber.

*N. benthamiana* plants were cultivated in a growth chamber with 16 h of light and 8 h of darkness at 22°C for 4 weeks. Subsequently, the 3rd or 4th leaf was detached and placed on moist tissue paper. Spores from *Phytophthora infestans* strain 88069 were harvested by addition of ddH_2_O to mycelium growing on plate. After 3-4 h incubation at 4°C, the zoospores were released by scratching the mycelium with a sterile tip. The zoospore concentration was adjusted to 10^5^ spores/ml of water.10µl of *P. infestans* spore suspension was dropped on detached leaves. After 6 days, the necrotic lesions were evaluated using ImageJ.

For *B. cinerea* infection assay, *B. cinerea B05*.*10* was grown 10 days on HA plates (10 g malt extract, 4 g glucose, 4 g yeast extract, 15 g agar per liter, pH 5.5). Spores were harvested and introduced at a concentration of 1x10^5^ conidia/ml to 10 ml Gamborg semi-solid inoculation medium at approcimately 37°C. The medium was incubated for 19h, 25°C, 200 rpm. 5 µl of the suspension was dropped on detached 6-week-old A. thaliana leaves (pretreated with *MbA* strains and *A. laibachii* as described in Eitzen et al. (2021). Lesion size was evaluated at 72hpi with ImageJ.

### Isolation of *A. laibachii* associated bacteria

*A. laibachii* spore suspension was prepared as described in in (Eitzen et al., 2021). After antibiotic treatment of the spores, spore solution was diluted 1:10, 1:100 and 1:1000 and 50 µl of each dilution and the undiluted suspension were plated on LB media. Plates were incubated at room temperature and after 3-5 days single colonies were picked. The identity of the isolated strains was determined via 16S rRNA sequencing using 16S rRNA primer pairs (**Table S1**). For each isolated bacterial strain, a colony PCR was performed with 16S rRNA primer pairs and purified PCR fragments were sequenced at the Eurofins sequencing facility (Germany). See **Table S2** for an overview of the isolated bacteria and the respective sequences.

### Microbial confrontation assays

For bacterial confrontation assays, *MbA* / *U. maydis* cultures grown to OD600nm 1.0 were dropped (10ul) in four quadrants of a PD Agar plate, spread previously with 100µl of bacterial culture with an OD600nm of 0.6. 20 µl of GH25 protein (0.8µg/µl) was applied to a hole created in the centre of the plate (d=4mm). For bacterial cross-inhibition assays, square PD plates were used. Bacteria were diluted to an OD600nm of 0.8 in 10 mM MgCl2 and 250 µl of a lawn bacterium was spread. 5 µl of competitor bacteria were dropped on the plate. Plates were incubated for 2-4 days at 22°C. For *P. infestans* confrontation assay, RSA plates supplemented with 100µg/ml Ampicillin were used. A mycelial plug of *P. infestans* was placed in the centre of the plate and growing culture of *MbA* (10µl) was placed as droplets on two corners of the plate. For *B. cinerea* confrontation assays, *MbA* strains were grown to an OD600:1.0 and 10 µl were dropped in the centre of an PD plate and incubated overnight at 22°C. 20 µl of MbA_GH25 (0.8µg/µl) was applied to a hole created in the centre of the plate (d=4mm). *B. cinerea B05*.*10* plugs were placed on two corners of the plate with the same distance to the competitor. Inhibition was evaluated at 3-4 dpi.

### Elicitor assays

2.5-week-old Col-0 *Arabidopsis thaliana* seedlings on liquid MS media were hand-infiltrated with purified recombinant proteins *MbA*_GH25, *MbA*_GH25(D124A), heat-killed *MbA*_GH25 in 2µM concentration. Bacterial PAMP, flg22 (50nM conc.) was used as a positive control. Leaves were harvested 30 minutes after treatment and frozen in liquid nitrogen. RNA and cDNA extracted from harvested tissue was used to perform qPCR of different marker genes. (modified from protocol of (Cao et al., 2014)).

### Nucleic acid methods

Total RNA extraction from plant samples were performed with Trizol Reagent (Invitrogen, Karlsruhe, Germany) according to the manufacturer’s instructions followed by treatment with Turbo DNA-Free Kit (Ambion/Applied Biosystems; Darmstadt, Germany) to remove any DNA contamination in the extracted RNA. Synthesis of cDNA was performed using First Strand cDNA Synthesis Kit (Thermo Fisher Scientific; Darmstadt, Germany) according to recommended instruction starting with a concentration of 10 µg RNA. RT-qPCR oligonucleotide pairs were designed with Primer3 Plus. Relative expression levels of marker genes were analyzed with GoTaq® qPCR Master Mix in a Bio-Rad iCycler system using the following program: 2 min at 95 °C followed by 45 cycles of 30 s at 95 °C, 30 s at 61 °C and 30 s 72 °C. Plasmid Prep Kit (QIAGEN, Venlo, The Netherlands) was used for isolation of plasmid DNA from bacteria after the principle of alkaline lysis. Genomic DNA was isolated using phenol–chloroform extraction protocol (Ruhe et al., 2016).

### Construction of plasmids

All plasmids were constructed using either restriction enzyme mediated ligation using T4 DNA ligase or Gibson assembly (New England Biolabs; Frankfurt a.M., Germany). All GH25 orthologue gene expression constructs (*p123-pUmOtef::MbA_GH25-2xHA, p123-pUmPit2:: MbA_GH25(D124A)2xHA*,*p123pUmOtef::Umaydis_GH252xHA, p123pUmOtef::Umaydis_G H25(D124A)-2xHA*,*p123pUmOtef::Rsolani_GH252xHA* and *p123pUmOtef::Rsolani_GH25(D122A)-2xHA)* were constructed using Gibson assembly. *Rhizoctonia solani* GH25 was di-codon optimized for expression in *U. maydis* and synthesized gene was obtained from BioCat GmbH, Heidelberg. GH25 active site mutants created by using QuickChange XL Site-Directed Mutagenesis Kit from Agilent Technology according to the manufacturer’s instructions. *E. coli* transformation was performed using heat shock according to standard molecular biology methods (Sambrook et al., 1989). Vector constructs and oligonucleotide sequences are mentioned in Table S1. Generated constructs were verified at the Eurofins sequencing facility (Germany).

### Fungal Transformation and Validation

The heterologous gene expression constructs (Table S1) were introduced in *SG200* or *FB1* (GH25 orthologue overexpression) by ip integration, using protoplasts according to (Kämper, 2004). Generated strains were further analyzed by southern blot to validate the integration and copy number (data not shown).

### Phylogenetic analysis

For phylogenetic tree construction, all selected GH25 orthologues (see Eitzen et al., (2021)) were aligned using MAFFT Multiple Sequence Alignment Software Version 7. Subsequently, a phylogenetic tree was constructed using NJ (Neighbor-Joining) with default settings and 100 bootstrap replications using Phyl.io (Robinson et al., 2016).

### Statistical analysis

For statistical analysis, to identify significant differences between treatments, an analysis of variance (ANOVA) model with Tukey’s HSD test was used for multiple comparisons. Graphpad Prism (10.4.1) was used to generate data plots. ImageJ 1.53K version (Wayne Rasband and Contributors National Institute of Health, USA) was used to calculate necrotic lesions of *P. infestans, B. cinerea* and zone of inhibition by fungal and yeast strains. Fiji (2.16.0) was used to analyse area of inhibition zones in bacterial confrontation assay. Di-Codon Optimization of *R. solani* GH25 was performed using Python 3.x (Python Software Foundation, 2023).

## Data availability

Source data are provided in this paper. All data supporting the findings of this study that are not directly available within the paper (and its supplementary data) will be upon reasonable request available from the corresponding authors (GD, PS).

## Acknowledgements

This project has received from the Cluster of Excellence on Plant Sciences (CEPLAS) funded under Germany’s Excellence Strategy—EXC 2048/1—project ID: 390686111 and the DFG priority program SPP2125 ‘DECRyPT’. We thank Prof. Laura Rose (HHU, Düsseldorf, Germany) and Prof. Francine Govers (Wageningen University, The Netherlands) for kindly providing the *P. infestans* strains.

## Author contributions

GD, ZS and PS designed the experiments with input from JP and EK. ZS, PS, KBH and JB conducted the experiments. PS, ZS and GD wrote the manuscript with contributions from all authors.

## Competing interests

The authors declare no competing interests.

## Supplementary Information

**Supplementary Figure S1:** Testing plant defense marker gene expression by MbA_GH25

**Supplementary Figure S2:** Interaction of A. laibachii cell wall with MbA_GH25

**Supplementary Figure S3:** Interaction of *MbA* and *MbA*_GH25 with oomycetes and fungi

**Supplementary Figure S4:** Inhibition of A. laibachii associated bacteria

**Supplementary Figure S5**: Inhibition of *A. laibachii* associated bacteria by GH25 orthologs

**Supplementary Figure S6**: Connection of cross-inhibition events *on-plate* and *A. laibachii* inhibition *on-planta*

**Supplementary Table S1**: Information on Constructs & Primer used

**Supplementary Table S2:** Information on bacterial strains used

## Notes

### Competing Interest Statement

The authors have declared no competing interest.

## References

Agler MT, Ruhe J, Kroll S, Morhenn C, Kim ST, Weigel D, Kemen EM. 2016. Microbial Hub Taxa Link Host and Abiotic Factors to Plant Microbiome Variation. PLoS Biol 14. doi:10.1371/JOURNAL.PBIO.1002352

Almario J, Mahmoudi M, Kroll S, Agler M, Placzek A, Mari A, Kemen E. 2022. The Leaf Microbiome of Arabidopsis Displays Reproducible Dynamics and Patterns throughout the Growing Season. mBio 13:e0282521. doi:10.1128/mbio.02825-21

Aronson JM, Cooper BA, Fuller MS. 1967. Glucans of Oomycete Cell Walls. Science (1979) 155:335– 336. doi:10.1126/SCIENCE.155.3760.332

Belhaj K, Cano LM, Prince DC, Kemen A, Yoshida K, Dagdas YF, Etherington GJ, Schoonbeek HJ, van Esse HP, Jones JDG, Kamoun S, Schornack S. 2017. Arabidopsis late blight: infection of a nonhost plant by Albugo laibachii enables full colonization by Phytophthora infestans. Cell Microbiol. doi:10.1111/cmi.12628

Berendsen RL, Vismans G, Yu K, Song Y, de Jonge R, Burgman WP, Burmølle M, Herschend J, Bakker PAHM, Pieterse CMJ. 2018. Disease-induced assemblage of a plant-beneficial bacterial consortium. ISME J 12:1496–1507. doi:10.1038/s41396-018-0093-1

Bradley EL, Ökmen B, Doehlemann G, Henrissat B, Bradshaw RE, Mesarich CH. 2022. Secreted Glycoside Hydrolase Proteins as Effectors and Invasion Patterns of Plant-Associated Fungi and Oomycetes. Front Plant Sci 13. doi:10.3389/FPLS.2022.853106

Brito N, Espino JJ, González C. 2007. The Endo-β-1,4-Xylanase Xyn11A Is Required for Virulence in Botrytis cinerea. https://doi.org/101094/MPMI-19-0025 19:p25–32. doi:10.1094/MPMI-19-0025

Cao Y, Liang Y, Tanaka K, Nguyen CT, Jedrzejczak RP, Joachimiak A, Stacey G. 2014. The kinase LYK5 is a major chitin receptor in Arabidopsis and forms a chitin-induced complex with related kinase CERK1. Elife 3:e03766. doi:10.7554/eLife.03766

Cernava T. 2024a. Coming of age for Microbiome gene breeding in plants. Nat Commun 15:6623. doi:10.1038/s41467-024-50700-7

Cernava T. 2024b. Coming of age for Microbiome gene breeding in plants. Nature Communications 2024 15:1 15:1–3. doi:10.1038/s41467-024-50700-7

Cooper AJ, Latunde-Dada AO, Woods-Tör A, Lynn J, Lucas JA, Crute IR, Holub EB. 2008. Basic compatibility of Albugo candida in Arabidopsis thaliana and Brassica juncea causes broad-spectrum suppression of innate immunity. Molecular Plant-Microbe Interactions. doi:10.1094/MPMI-21-6-0745

de Angelis G, Simonetti G, Chronopoulou L, Orekhova A, Badiali C, Petruccelli V, Portoghesi F, D’Angeli S, Brasili E, Pasqua G, Palocci C. 2022. A novel approach to control Botrytis cinerea fungal infections: uptake and biological activity of antifungals encapsulated in nanoparticle based vectors. Sci Rep 12:7989. doi:10.1038/s41598-022-11533-w

de Vries S, von Dahlen JK, Schnake A, Ginschel S, Schulz B, Rose LE. 2018. Broad-spectrum inhibition of Phytophthora infestans by fungal endophytes. FEMS Microbiol Ecol 94. doi:10.1093/FEMSEC/FIY037

de Vrieze M, Gloor R, Codina JM, Torriani S, Gindro K, L’Haridon F, Bailly A, Weisskopf L. 2019. Biocontrol Activity of Three Pseudomonas in a Newly Assembled Collection of Phytophthora infestans Isolates. Phytopathology 109:1555–1565. doi:10.1094/PHYTO-12-18-0487-R

Eitzen K, Sengupta P, Kroll S, Kemen E, Doehlemann G. 2021. A fungal member of the Arabidopsis thaliana phyllosphere antagonizes Albugo laibachii via a GH25 lysozyme. Elife 10:e65306. doi:10.7554/eLife.65306

Flores-Nunez VM, Stukenbrock EH. 2024. The impact of filamentous plant pathogens on the host microbiota. BMC Biol 22:175. doi:10.1186/s12915-024-01965-3

Freimoser FM, Rueda-Mejia MP, Tilocca B, Migheli Q. 2019. Biocontrol yeasts: mechanisms and applications. World Journal of Microbiology and Biotechnology 2019 35:10 35:1–19. doi:10.1007/S11274-019-2728-4

Gfeller A, Fuchsmann P, De Vrieze M, Gindro K, Weisskopf L. 2022. Bacterial Volatiles Known to Inhibit Phytophthora infestans Are Emitted on Potato Leaves by Pseudomonas Strains. Microorganisms 2022, Vol 10, Page 1510 10:1510. doi:10.3390/MICROORGANISMS10081510

Gómez-Pérez D, Schmid M, Chaudhry V, Hu Y, Velic A, Maček B, Ruhe J, Kemen A, Kemen E. 2023. Proteins released into the plant apoplast by the obligate parasitic protist Albugo selectively repress phyllosphere-associated bacteria. New Phytologist 239:2320–2334. doi:10.1111/nph.18995

Goossens P, Spooren J, Baremans KCM, Andel A, Lapin D, Echobardo N, Pieterse CMJ, Van den Ackerveken G, Berendsen RL. 2023. Obligate biotroph downy mildew consistently induces near-identical protective microbiomes in Arabidopsis thaliana. Nat Microbiol 8:2349–2364. doi:10.1038/s41564-023-01502-y

Gui YJ, Chen JY, Zhang DD, Li NY, Li TG, Zhang WQ, Wang XY, Short DPG, Li L, Guo W, Kong ZQ, Bao YM, Subbarao K V., Dai XF. 2017. Verticillium dahliae manipulates plant immunity by glycoside hydrolase 12 proteins in conjunction with carbohydrate-binding module 1. Environ Microbiol 19:1914–1932. doi:10.1111/1462-2920.13695

Henrissat B, Davies G. 1997. Structural and sequence-based classification of glycoside hydrolases. Curr Opin Struct Biol 7:637–644. doi:10.1016/S0959-440X(97)80072-3

Jurburg SD, Eisenhauer N, Buscot F, Chatzinotas A, Chaudhari NM, Heintz-Buschart A, Kallies R, Küsel K, Litchman E, Macdonald CA, Müller S, Reuben RC, da Rocha UN, Panagiotou G, Rillig MC, Singh BK. 2022. Potential of microbiome-based solutions for agrifood systems. Nat Food 3:557–560. doi:10.1038/s43016-022-00576-x

Kamoun S, Furzer O, Jones JDG, Judelson HS, Ali GS, Dalio RJD, Roy SG, Schena L, Zambounis A, Panabières F, Cahill D, Ruocco M, Figueiredo A, Chen X-R, Hulvey J, Stam R, Lamour K, Gijzen M, Tyler BM, Grünwald NJ, Mukhtar MS, Tomé DFA, Tör M, Van Den Ackerveken G, McDowell J, Daayf F, Fry WE, Lindqvist-Kreuze H, Meijer HJG, Petre B, Ristaino J, Yoshida K, Birch PRJ, Govers F. 2015. The Top 10 oomycete pathogens in molecular plant pathology. Mol Plant Pathol 16:413–434. doi:10.1111/mpp.12190

Kämper J. 2004. A PCR-based system for highly efficient generation of gene replacement mutants in Ustilago maydis. Molecular Genetics and Genomics. doi:10.1007/s00438-003-0962-8

Kettles GJ, Bayon C, Sparks CA, Canning G, Kanyuka K, Rudd JJ. 2018. Characterization of an antimicrobial and phytotoxic ribonuclease secreted by the fungal wheat pathogen Zymoseptoria tritici. New Phytologist 217:320–331. doi:10.1111/nph.14786

Köhl J, Kolnaar R, Ravensberg WJ. 2019. Mode of Action of Microbial Biological Control Agents Against Plant Diseases: Relevance Beyond Efficacy. Front Plant Sci 10. doi:10.3389/fpls.2019.00845

Kong P, Hong C. 2016. Soil bacteria as sources of virulence signal providers promoting plant infection by Phytophthora pathogens. Scientific Reports 2016 6:1 6:1–13. doi:10.1038/srep33239

Loc NH, Huy ND, Quang HT, Lan TT, Thu Ha TT. 2020. Characterisation and antifungal activity of extracellular chitinase from a biocontrol fungus, Trichoderma asperellum PQ34. Mycology 11:38– 48. doi:10.1080/21501203.2019.1703839

Ma Z, Song T, Zhu L, Ye W, Wang Yang, Shao Y, Dong S, Zhang Z, Dou D, Zheng X, Tyler BM, Wang Yuanchao. 2015. A Phytophthora sojae Glycoside Hydrolase 12 Protein Is a Major Virulence Factor during Soybean Infection and Is Recognized as a PAMP. Plant Cell 27:2057–2072. doi:10.1105/TPC.15.00390

Mahmoudi M, Almario J, Hu Y, Tenzer L-M, Nieselt K, Kemen E. 2024. Biotic interactions shape infection outcomes in Arabidopsis. bioRxiv 2024.10.25.620230. doi:10.1101/2024.10.25.620230

Mélida H, Sandoval-Sierra J V., Diéguez-Uribeondo J, Bulone V. 2013. Analyses of Extracellular Carbohydrates in Oomycetes Unveil the Existence of Three Different Cell Wall Types. Eukaryot Cell 12:194. doi:10.1128/EC.00288-12

Morita T, Konishi M, Fukuoka T, Imura T, Kitamoto D. 2007. Microbial conversion of glycerol into glycolipid biosurfactants, mannosylerythritol lipids, by a basidiomycete yeast, Pseudozyma antarctica JCM 10317T. J Biosci Bioeng. doi:10.1263/jbb.104.78

Ökmen B, Katzy P, Huang L, Wemhöner R, Doehlemann G. 2023. A conserved extracellular Ribo1 with broad-spectrum cytotoxic activity enables smut fungi to compete with host-associated bacteria. New Phytologist 240:1976–1989. doi:10.1111/nph.19244

Ordon J, Gantner J, Kemna J, Schwalgun L, Reschke M, Streubel J, Boch J, Stuttmann J. 2017. Generation of chromosomal deletions in dicotyledonous plants employing a user-friendly genome editing toolkit. The Plant Journal 89:155–168. doi:10.1111/tpj.13319

Oubaha B, Ezzanad A, Bolívar-Anillo HJ, Oubaha B, Ezzanad A, Bolívar-Anillo HJ. 2021. Plant Beneficial Microbes Controlling Late Blight Pathogen Phytophthora infestans. Agro-Economic Risks of Phytophthora and an Effective Biocontrol Approach. doi:10.5772/INTECHOPEN.99383

Prince DC, Rallapalli G, Xu D, Schoonbeek H, Çevik V, Asai S, Kemen E, Cruz-Mireles N, Kemen A, Belhaj K, Schornack S, Kamoun S, Holub EB, Halkier BA, Jones JDG. 2017. Albugo-imposed changes to tryptophan-derived antimicrobial metabolite biosynthesis may contribute to suppression of non-host resistance to Phytophthora infestans in Arabidopsis thaliana. BMC Biol 15:20. doi:10.1186/s12915-017-0360-z

Rau A, Hogg T, Marquardt R, Hilgenfeld R. 2001. A new lysozyme fold. Crystal structure of the muramidase from Streptomyces coelicolor at 1.65 A resolution. J Biol Chem 276:31994–31999. doi:10.1074/JBC.M102591200

Robinson O, Dylus D, Dessimoz C. 2016. Phylo.io: Interactive Viewing and Comparison of Large Phylogenetic Trees on the Web. Mol Biol Evol 33:2163–2166. doi:10.1093/molbev/msw080

Ruhe J, Agler MT, Placzek A, Kramer K, Finkemeier I, Kemen EM. 2016. Obligate biotroph pathogens of the genus Albugo are better adapted to active host defense compared to niche competitors. Front Plant Sci 7:184119. doi:10.3389/FPLS.2016.00820/BIBTEX

Sambrook J, Fritsch EF, Maniatis T. 1989. Molecular cloning: a laboratory manual. Cold Spring Harbor, NY: Cold Spring Harbor Laboratory Press.

Saravanakumar K, Dou K, Lu Z, Wang X, Li Y, Chen J. 2018. Enhanced biocontrol activity of cellulase from Trichoderma harzianum against Fusarium graminearum through activation of defense-related genes in maize. Physiol Mol Plant Pathol 103:130–136. doi:10.1016/j.pmpp.2018.05.004

Singh BK, Delgado-Baquerizo M, Egidi E, Guirado E, Leach JE, Liu H, Trivedi P. 2023. Climate change impacts on plant pathogens, food security and paths forward. Nat Rev Microbiol 21:640– 656. doi:10.1038/s41579-023-00900-7

Snelders NC, Kettles GJ, Rudd JJ, Thomma BPHJ. 2018. Plant pathogen effector proteins as manipulators of host microbiomes? Mol Plant Pathol 19:257–259. doi:10.1111/MPP.12628

Snelders NC, Rovenich H, Thomma BPHJ. 2022. Microbiota manipulation through the secretion of effector proteins is fundamental to the wealth of lifestyles in the fungal kingdom. FEMS Microbiol Rev 46:fuac022. doi:10.1093/femsre/fuac022

Thines M. 2014. Phylogeny and evolution of plant pathogenic oomycetes-a global overview. Eur J Plant Pathol. doi:10.1007/s10658-013-0366-5

Tzelepis G, Dubey M, Jensen DF, Karlsson M. 2015. Identifying glycoside hydrolase family 18 genes in the mycoparasitic fungal species Clonostachys rosea. Microbiology (N Y) 161:1407–1419. doi:10.1099/mic.0.000096

van der Biezen EA, Dreddie CT, Kahn K, Parker JE, Jones JDG. 2002. Arabidopsis RPP4 is a member of the RPP5 multigene family of TIR-NB-LRR genes and confers downy mildew resistance through multiple signalling components. The Plant Journal, 29:439–451. 10.1046/j.0960-7412.2001.01229.x

Weiberg A, Wang M, Lin FM, Zhao H, Zhang Z, Kaloshian I, Huang H Da, Jin H. 2013. Fungal small RNAs suppress plant immunity by hijacking host RNA interference pathways. Science (1979) 342:118–123. doi:10.1126/SCIENCE.1239705

Yang L, Qu M, Wang Z, Huang S, Wang Q, Wei M, Li F, Yang D, Pan L. 2024. Biochemical Properties of a Novel Cold-Adapted GH19 Chitinase with Three Chitin-Binding Domains from Chitinilyticum aquatile CSC-1 and Its Potential in Biocontrol of Plant Pathogenic Fungi. J Agric Food Chem 72:19581–19593. doi:10.1021/acs.jafc.4c02559

Yu C, Li T, Shi X, Saleem M, Li B, Liang W, Wang C. 2018. Deletion of endo-β-1,4-xylanase VmXyl1 impacts the virulence of Valsa mali in apple tree. Front Plant Sci 9. doi:10.3389/FPLS.2018.00663/FULL

Yu C, Liang X, Song Y, Ali Q, Yang X, Zhu L, Gu Q, Kuptsov V, Kolomiets E, Wu H, Gao X. 2024. A glycoside hydrolase 30 protein BpXynC of Bacillus paralicheniformis NMSW12 recognized as A MAMP triggers plant immunity response. Int J Biol Macromol 261:129750. doi:10.1016/j.ijbiomac.2024.129750

Zhang L, Yan J, Fu Z, Shi W, Ninkuu V, Li G, Yang X, Zeng H. 2021. FoEG1, a secreted glycoside hydrolase family 12 protein from Fusarium oxysporum, triggers cell death and modulates plant immunity. Mol Plant Pathol 22:522–538. doi:10.1111/MPP.13041

