## Supplemental Figures for "GH25 lysozyme mediates tripartite interkingdom interactions and microbial competition on the plant leaf surface"

* Contributed equally to the work

#Authors for correspondence:

### Supplementary Figures

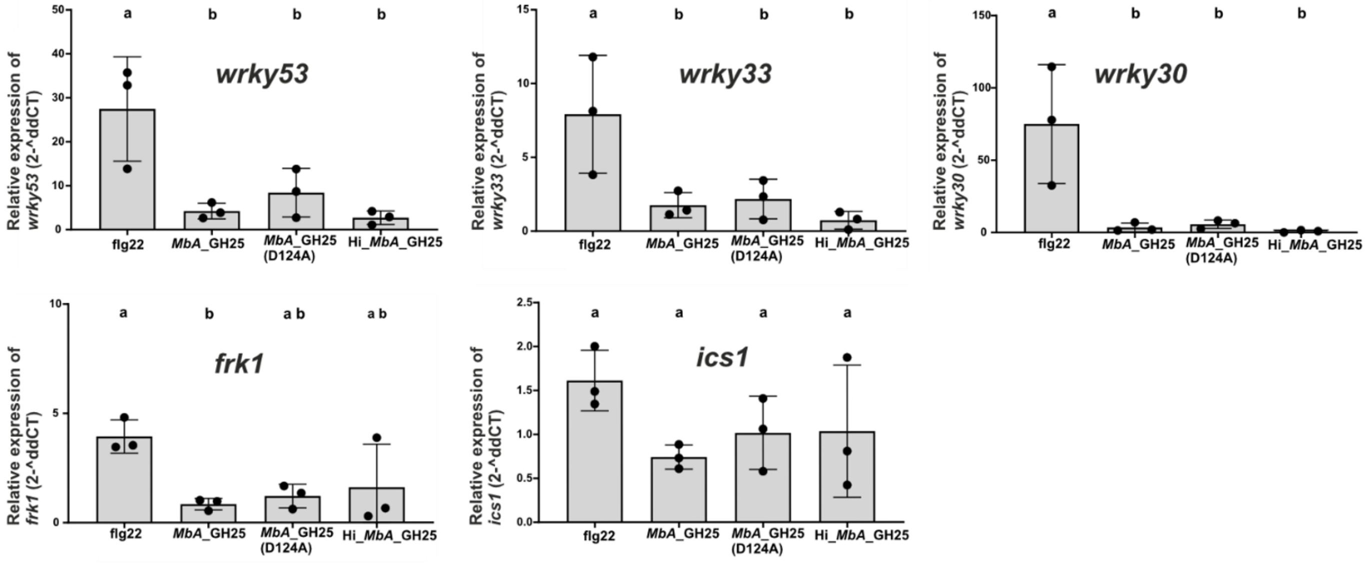


**Figure S1: Testing plant defense marker gene expression by *MbA*_GH25.** Expression levels of marker genes *WRKY30, WRKY33, WRKY53, FRK 1*, and *ICS1* in *Arabidopsis thaliana* (Col-0 ecotype) relative to the mock treatment (1/2 liquid MS), normalized to ef1α. Each data point corresponds to one biological replicate. Error bars indicate SD. One-way ANOVA and Tukey's multiple comparisons test (alpha 0.05) was performed to find significant difference between treatments, indicated by letters above.

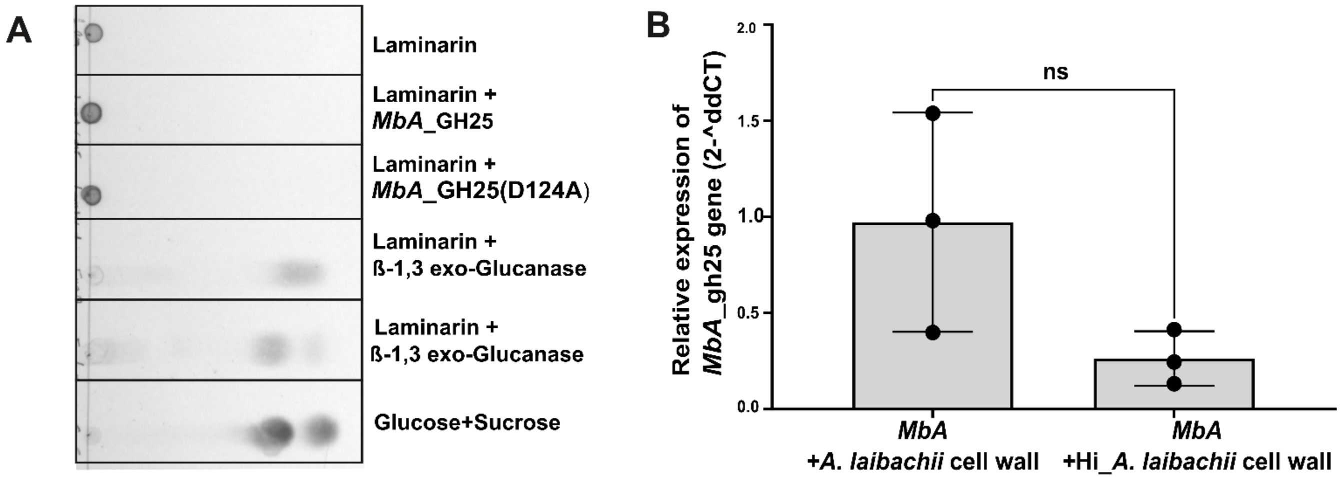


**Figure S2: Interaction of *A. laibachii* cell wall with *MbA*_GH25**. **A)** Thin layer Chromatography of Laminarin: Samples were loaded on TLC Silica gel 60 F254 plate. Glucose + Sucrose mix was used as reference. Laminarin and their hydrolysis products were visualized by spraying the TLC plate with detection solution. Hydrolysis products were observed for the Laminarin treated with ß-1,3 exo and endoglucanase, but not with *MbA*_GH25 (active and mutated) protein. **B)** Relative *MbA*_gh25 gene expression analysis in *MbA* liquid culture in presence of *A. laibachii* cell wall (active and heat-inactivated-Hi), normalized to *MbA* culture + buffer control using *MbA*_ppi housekeeping genes in ddCT method. Unpaired t-test with p value=0.1. Error bar indicates SD. No significant difference observed between treatments.

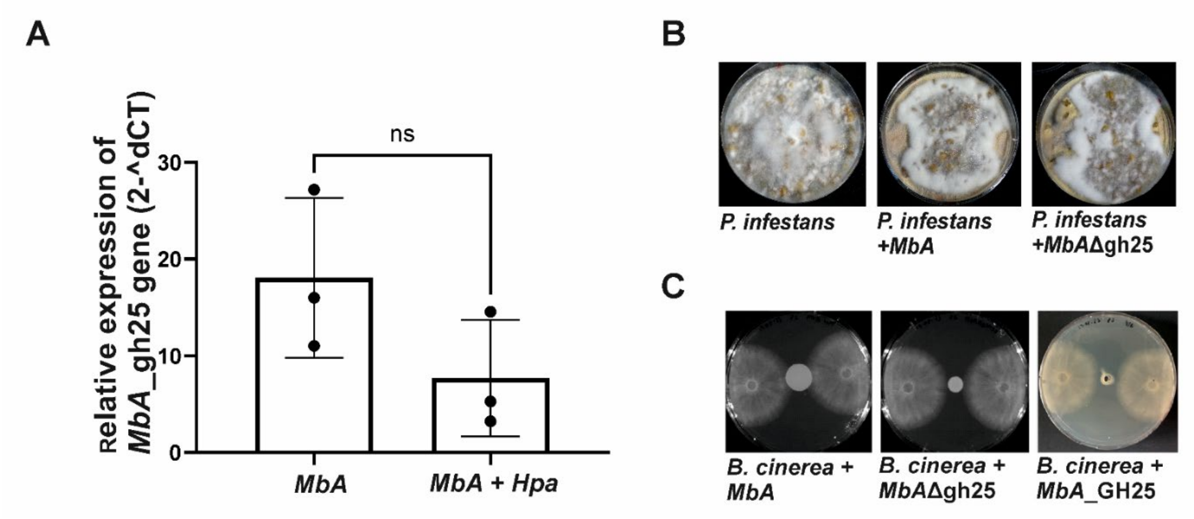


**Figure S3: Interaction of *MbA* and *MbA*_GH25 with oomycetes and fungi. A)**-Relative gh25 gene expression analysis in *MbA* in response to *Hpa* treatment in *A. thaliana* Col-0, normalized to *MbA* housekeeping gene ppi (unpaired t-test, P=0.15). **B)**-Confrontation assay between *MbA*/*MbA*Δgh25 and *P. infestans* strain: *in vitro* on RSA plate where no zone of inhibition is observed. **C)**-plate-based inhibition assay showed no inhibition of *B. cinerea* against *MbA*, *MbA*Δgh25and purified *MbA*_GH25. Strains were dropped at the centre of the plate (10 µl of culture with OD600nm = 1.0). 20 µl of *MbA*_GH25 (0.8 mg/ml) was applied to hole on the centre of the plate. Two agar discs with *B. cinerea* spores were placed on opposing sides of the plate and growth around the competitor was observed at 3 dpi.

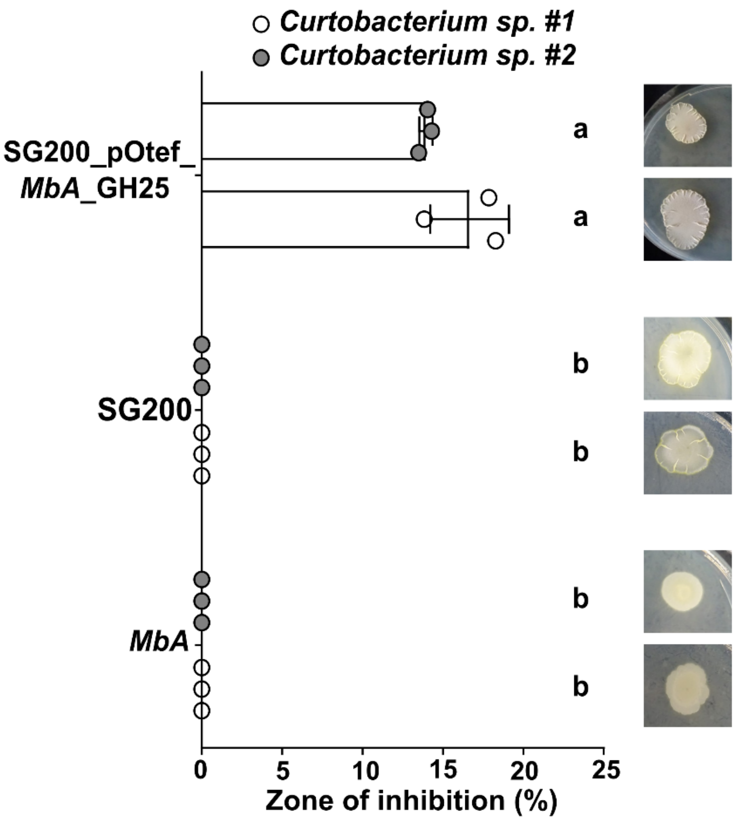


**Figure S4: Inhibition of *A. laibachii* associated bacteria.** % zone of inhibition (halo formation) at 3 dpi was calculated for the strains *Pseduomonas* sp. (inhibited by *MbA*_WT) and *Curtobacterium* sp. (inhibited by (SG200_pOtef:*MbA*GH25). *U. maydis* strain SG200 used as a negative control which did not show any inhibition towards any bacteria across 3 independent replicates. Representative image of bacterial inhibition was added next to each treatment as indicated by halo formation surrounding the single colony of yeast/fungi. Error bars show SD. One-way ANOVA and Tukey's multiple comparisons test (alpha 0.05) was performed to find significant difference between treatments.

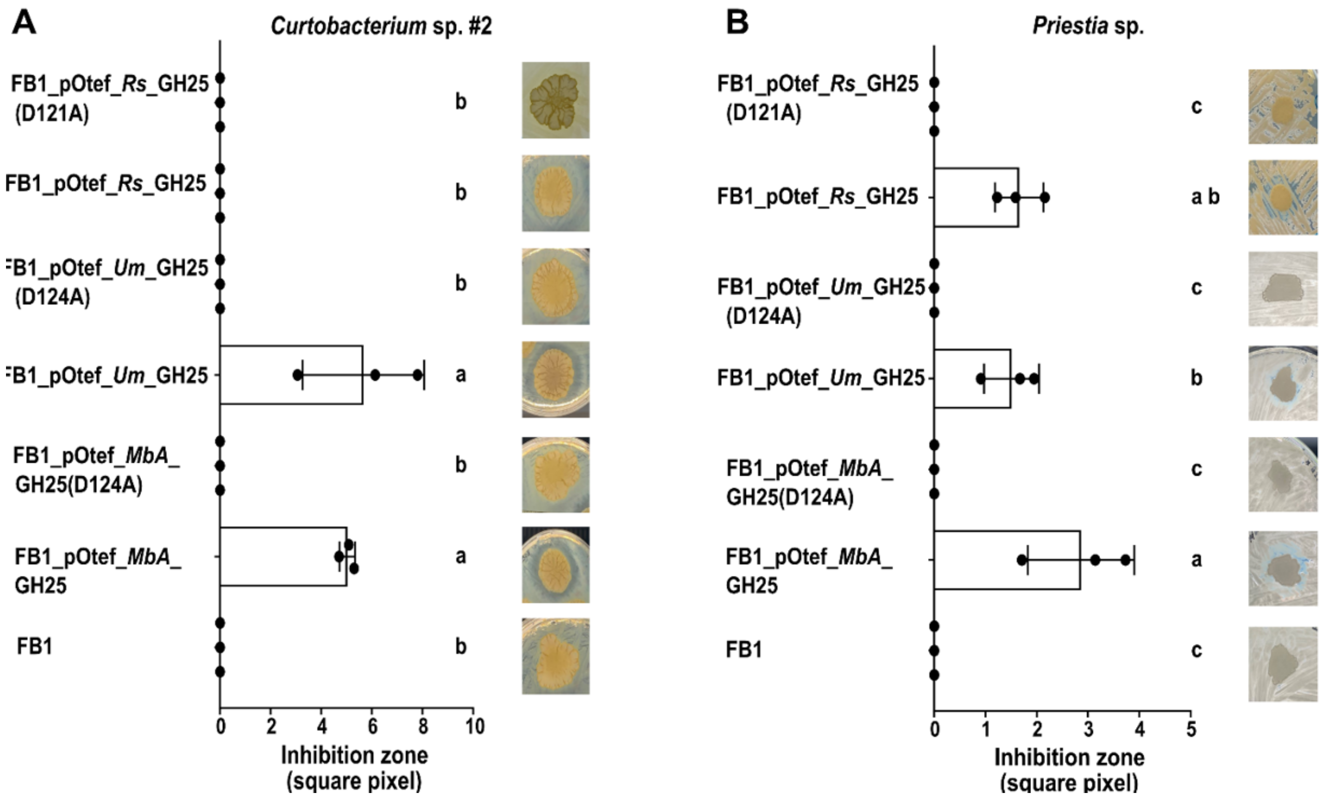


**Figure S5: Inhibition of *A. laibachii* associated bacteria by GH25 orthologs.** FB1 strain and recombinant FB1 strain overexpressing GH25 orthologues from *MbA*, *U. maydis* (*Um*) and *R. solani* (*Rs*) in active or mutant version were tested in a confrontation assay with **A)** *Curtobacterium* sp. #2 and **B)** *Priestia* sp. The inhibition zones surrounding fungal colonies were plotted across 3 independent replicates. One-way ANOVA and Tukey's multiple comparisons test (alpha 0.05) was performed to find significant difference between treatments. Representative image of bacterial inhibition was added next to each treatment as indicated by halo formation surrounding the single colony of yeast/fungi. Error bars show SD.

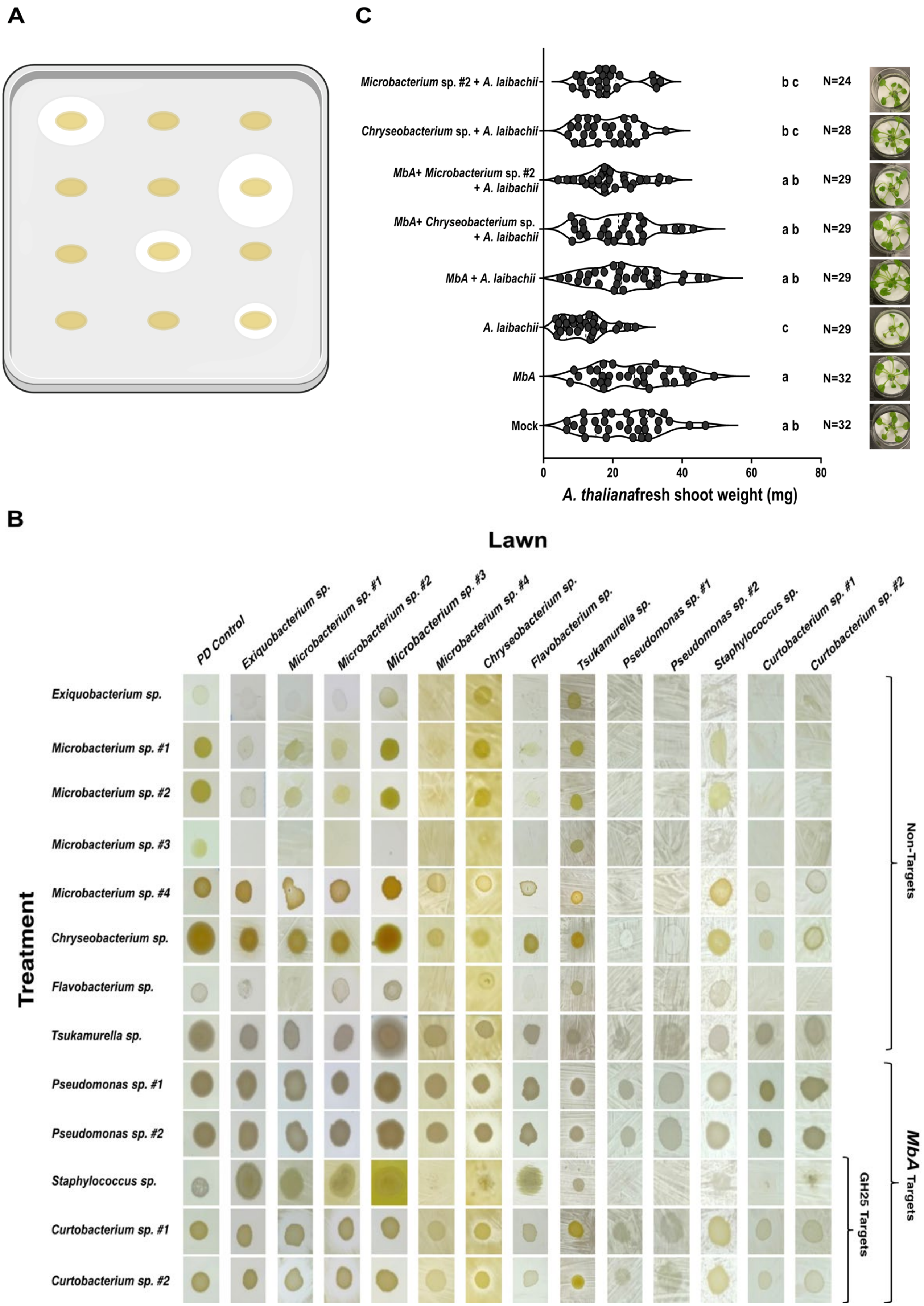


**Figure S6: Connection of cross-inhibition events *on-plate* and *A. laibachii* inhibition *on-planta*. A)** Schematic overview of experimental setup and B) Representative pictures of growth phenotype of each treatment bacterium spotted on different lawn bacteria. *Pseudomonas* sp. and *Curtobacterium* sp. lead to a strong growth reduction of most treated non-target bacteria. All isolated bacteria from *A. laibachii* spores were diluted to an OD600nm of 0.8 in 10 mM MgCl2 and 250 µl of the bacterial suspension was plated on square PD plates as lawn. 5 µl of each bacterial suspension was spottet on top as treatments. Location of the droplet was altered between replicates; 3-4 biological replicates were performed for each bacterial competition pair tested. **C.)** Fresh shoot weight measured at 11dpi showed *A. thaliana* seedlings to be reduced in growth upon infection with *A. laibachii* compared to mock treated seedlings. Pre-treatment with *MbA* with and without *Microbacterium* sp. #2 and *Chryseobacterium* sp. led to a higher recovery of fresh shoot weight in *A. laibachii* infected seedlings. Moreover, also *Microbacterium* sp. #2 and *Chryseobacterium* sp. treatment alone could recover fresh shoot weight in *A. laibachii* infected seedlings. The number of seedlings (N) were measured across 3 biological replicates. One-way ANOVA and Tukey' HSD (multiple comparisons of means; 95% family-wise confidence level) was performed to find significant difference between treatments. Representative image of *A. thaliana* seedlings is added next to each treatment.
